# Complete connectomic reconstruction of olfactory projection neurons in the fly brain

**DOI:** 10.1101/2020.01.19.911453

**Authors:** A.S. Bates, P. Schlegel, R.J.V. Roberts, N. Drummond, I.F.M. Tamimi, R. Turnbull, X. Zhao, E.C. Marin, P.D. Popovici, S. Dhawan, A. Jamasb, A. Javier, F. Li, G.M. Rubin, S. Waddell, D.D. Bock, M. Costa, G.S.X.E. Jefferis

**Author notes:** These authors contributed equally to this work (A.S. Bates, P. Schlegel).

## Abstract

Nervous systems contain sensory neurons, local neurons, projection neurons and motor neurons. To understand how these building blocks form whole circuits, we must distil these broad classes into neuronal cell types and describe their network connectivity. Using an electron micrograph dataset for an entire *Drosophila melanogaster* brain, we reconstruct the first complete inventory of olfactory projections connecting the antennal lobe, the insect analogue of the mammalian olfactory bulb, to higher-order brain regions in an adult animal brain. We then connect this inventory to extant data in the literature, providing synaptic-resolution ‘holotypes’ both for heavily investigated and previously unknown cell types. Projection neurons are approximately twice as numerous as reported by light level studies; cell types are stereotyped, but not identical, in cell and synapse numbers between brain hemispheres. The lateral horn, the insect analogue of the mammalian cortical amygdala, is the main target for this olfactory information and has been shown to guide innate behaviour. Here, we find new connectivity motifs, including: axo-axonic connectivity between projection neurons; feedback and lateral inhibition of these axons by local neurons; and the convergence of different inputs, including non-olfactory inputs and memory-related feedback onto lateral horn neurons. This differs from the configuration of the second most prominent target for olfactory projection neurons: the mushroom body calyx, the insect analogue of the mammalian piriform cortex and a centre for associative memory. Our work provides a complete neuroanatomical platform for future studies of the adult *Drosophila* olfactory system.

**Highlights:**

- **First complete parts list for second-order neurons of an adult olfactory system**
- **Quantification of left-right stereotypy in cell and synapse number**
- **Axo-axonic connections form hierarchical communities in the lateral horn**
- **Local neurons and memory-related feedback target projection neuron axons**

## 1 Introduction

Peripheral sensory neurons comprising the first layer of the olfactory system encode information about an animal’s chemical environment. Central neurons must then extract salient features from this information and use them to guide the animal’s behaviour. This guidance also depends on cocomputation with other sensory modalities as well as internal states, especially memory, with which olfaction is strongly linked. We now have a good mechanistic understanding of the principles of information processing in the primary olfactory centre where first and second-order neurons meet; this logic appears very similar between insects and mammals [1, 2]. However, even in the intensively studied vinegar fly, *Drosophila melanogaster*, much less is known about what happens in olfactory circuits beyond the second layer [3, 4].

In the insect brain, olfactory information is first received in the antennal lobe (AL), analogous to the mammalian olfactory bulb. Here, the axons of first-order olfactory sensory neurons (OSNs) ramify in 51 distinct olfactory glomeruli, a figure we finalise in the present work. Each glomerulus is the target of a specific type of OSN defined by its odorant receptor(s). In addition to these olfactory glomeruli, there are 7 thermo- and hygrosensory glomeruli in the ventral-posterior (VP) part of the AL (Marin et al., in prep.). Within the AL various local computations take place, e.g. divisive normalisation via lateral inhibition, before projection neurons (PNs), analogous to vertebrate mitral/tufted cells, carry information to higher-order brain regions (Figure 1A-B) [2]. PNs of the same cell type display broadly similar odour tuning across animals [5–8].

**Figure 1:**
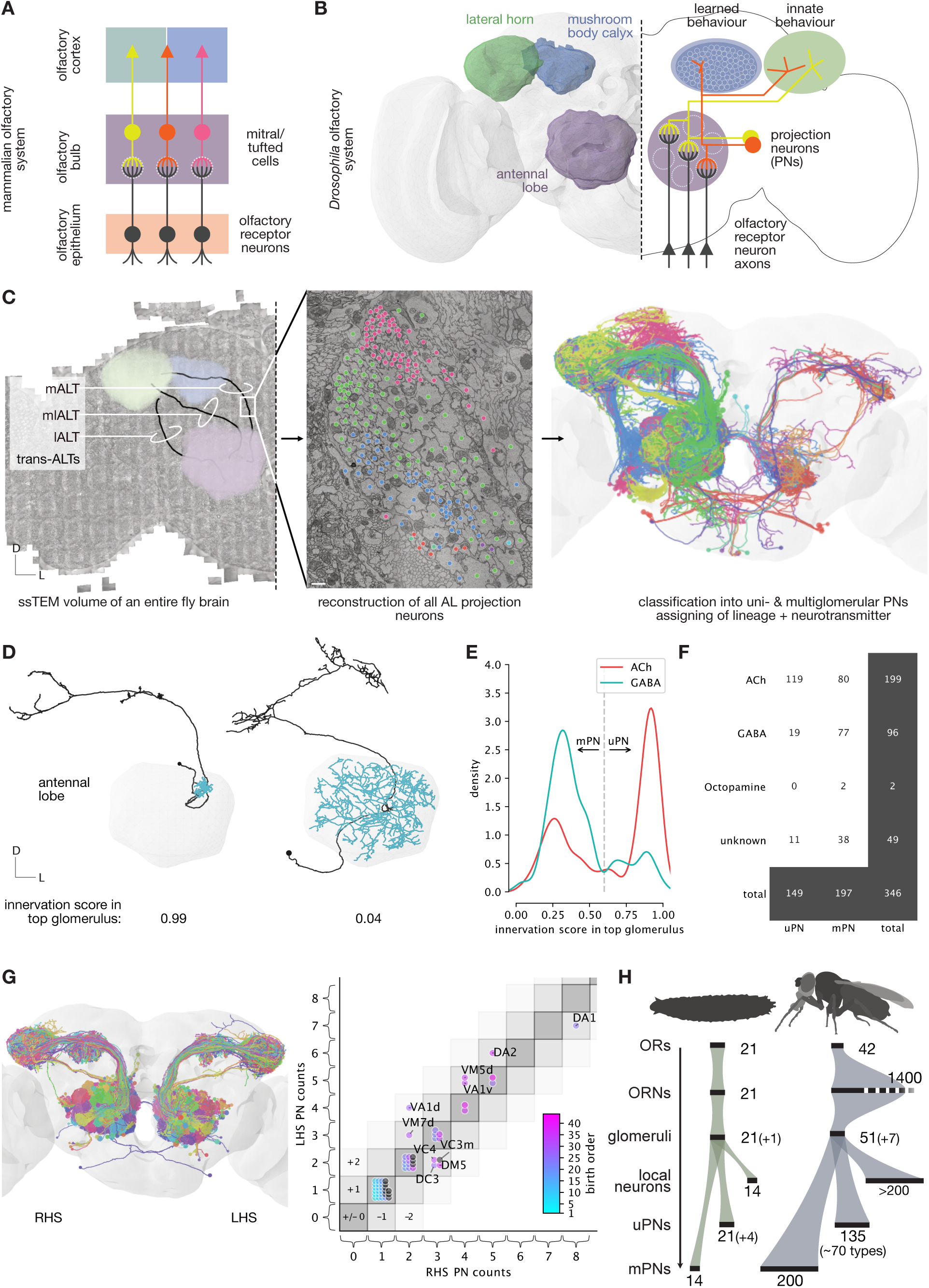
Second-order olfactory projection neurons. **A** Layout of the mammalian olfactory system. **B** The olfactory system in fruit flies shares similarities with that of mammals. **C** Second-order olfactory projection neurons (PNs) in all antennal lobe tracts (ALT) were reconstructed from a serial section transmission electron microscopy (ssTEM) volume of an entire fly brain. Scale bar represents 1 micron. **D** Exemplary sparse (left) and broad (right) PN. Dendrites used for classification in blue. **E** PNs were classified as uni- (uPN) or multiglomerular (mPN) projection neurons based on the sparseness of their glomerular innervation (see also Figure S1). **F** PN counts by class and neurotransmitter. **G** Comparison of right- (RHS) vs left-hand-side (LHS) cell counts for 58 of the 78 uPN types. **H** Scaling of the olfactory system from larval to adult *D. melanogaster*.

Olfactory PNs target multiple brain regions, including the well-studied insect mushroom body (MB), a structure analogous to the piriform cortex in mammals as both areas enable associative memory with respect to odours [9–12] (Figure 1B). Recent work indicates that higher-order olfactory systems are extensively interconnected and that a proper understanding of olfactory memory must emerge from an integrated understanding across multiple olfactory neuropils [13–15]. However, the organisation of other third-order olfactory areas that instigate innate behaviours is not well understood.

Headway has been made in elucidating architectural and functional aspects across higher-order, insect olfactory circuits [5, 13, 15–20]. Currently, we lack a framework for the olfactory system to help contextualise these results. Such a framework would need to consist of a complete and detailed description of the constituent blocks (i.e. cell types) and connections comprising the circuitry. We now provide a full inventory of olfactory inputs to higher brain centres and use this to shed light on third-order olfactory processing. This work creates a neuroinformatic scaffold that can be related to diverse data from other studies.

## Results

Electron microscopy (EM) is the only means by which to resolve fine neurites (<200 nm), synaptic vesicles (∼ 40 nm) and synaptic clefts (∼ 20 nm) and so see all synaptic connections between definable neurons in a sample. In order to create a full inventory of projection neurons (PNs) from the antennal lobe (AL), we turned to a recent EM dataset for a Full Adult Fly Brain (FAFB) [21].

### A full account of olfactory projection neurons connecting second and third-order brain areas

Previously published PN surveys were based on sparse labelling and therefore likely incomplete. We found 346 neurons with dendrites in the right AL and axonal projections to other neuropils (Figure 1C, see Methods). We cross-referenced these PNs with extant data [22, 23], linking neurons to their developmental hemilineages and putative neurotransmitter expression (Figure S1A; Supplemental Data; see Methods) [24–28].

PN dendrites exhibit diverse morphologies ranging from targeting only a single glomerulus to covering almost the entire AL (Figure 1D). To quantitatively describe their AL innervation patterns, we calculated a PN by glomerulus innervation matrix (Figure S1B, see Methods). This enabled us to calculate sparseness as a function of how many glomeruli are targeted by a given a PN; cholinergic PNs appear, on average, to be sparser than GABAergic PNs (Figure 1E).

There is no apparent threshold that distinguishes uniglomerular (uPNs) from multiglomerular PNs (mPNs). We, therefore, used well-studied uPNs [21, 23, 29–31] to set a sparseness threshold and classify PNs as either uni- or multiglomerular (Figure 1B). Recent work based on the same EM dataset lists 114 uPNs [13, 21]. We now update this count to 149 uni- and 197 multiglomerular PNs; 2-3 times more than previously reported [23, 29, 30]. Most uPNs are cholinergic while mPNs are about evenly split between cholinergic and GABAergic (Figure 1F).

We define 78 uPN types based on their (hemi-) lineage, the glomerulus innervated and the antennal lobe tract they project through (see Supplemental Data). To assess numerical variability, we reconstructed all members of 58 of these types on the left-hand-side to identification and matched them to their contralateral homologs. We find left/right numbers to vary by 1.1 ±0.3 s.d. for 10 out of the 58 tested uPN types (Figure 1G).

Larval and adult uPNs have previously been shown to receive inputs from a variety of sources: olfactory sensory neurons (OSNs), local neurons and other PNs [2, 32–34]. We examined a single mPN (Figure S1C) and found that direct OSN input is present in a subset of glomeruli. Input in other glomeruli comes exclusively from local neurons and other PNs. Odour responses of mPNs might therefore be more complex than simple summation across odour channels; mPNs could receive feedforward excitation in glomerulus A but pick up lateral inhibition in glomeruli B.

### The synaptic organisation of olfactory projection neurons

We next turned to third-order olfactory neuropils and, for each PN, reconstructed its ipsilateral axon to synaptic completion (Figure 2A and S2A, see Methods). The lateral horn (LH) and the Calyx (CA) are the main second-order targets for olfactory information, both in terms of number of PNs targeting those neuropils as well as the total number of PN presynapses placed in either neuropil (Figure 2B,C and S2B). However, the LH differs from the CA in that it receives almost all the AL’s feedforward inhibition. Moreover, PN axons in the LH, but not the CA, are heavily modulated. For example, the uPN for glomerulus DP1m receives 1236 synaptic inputs onto its axon in the LH but only 50 in the CA.

**Figure 2:**
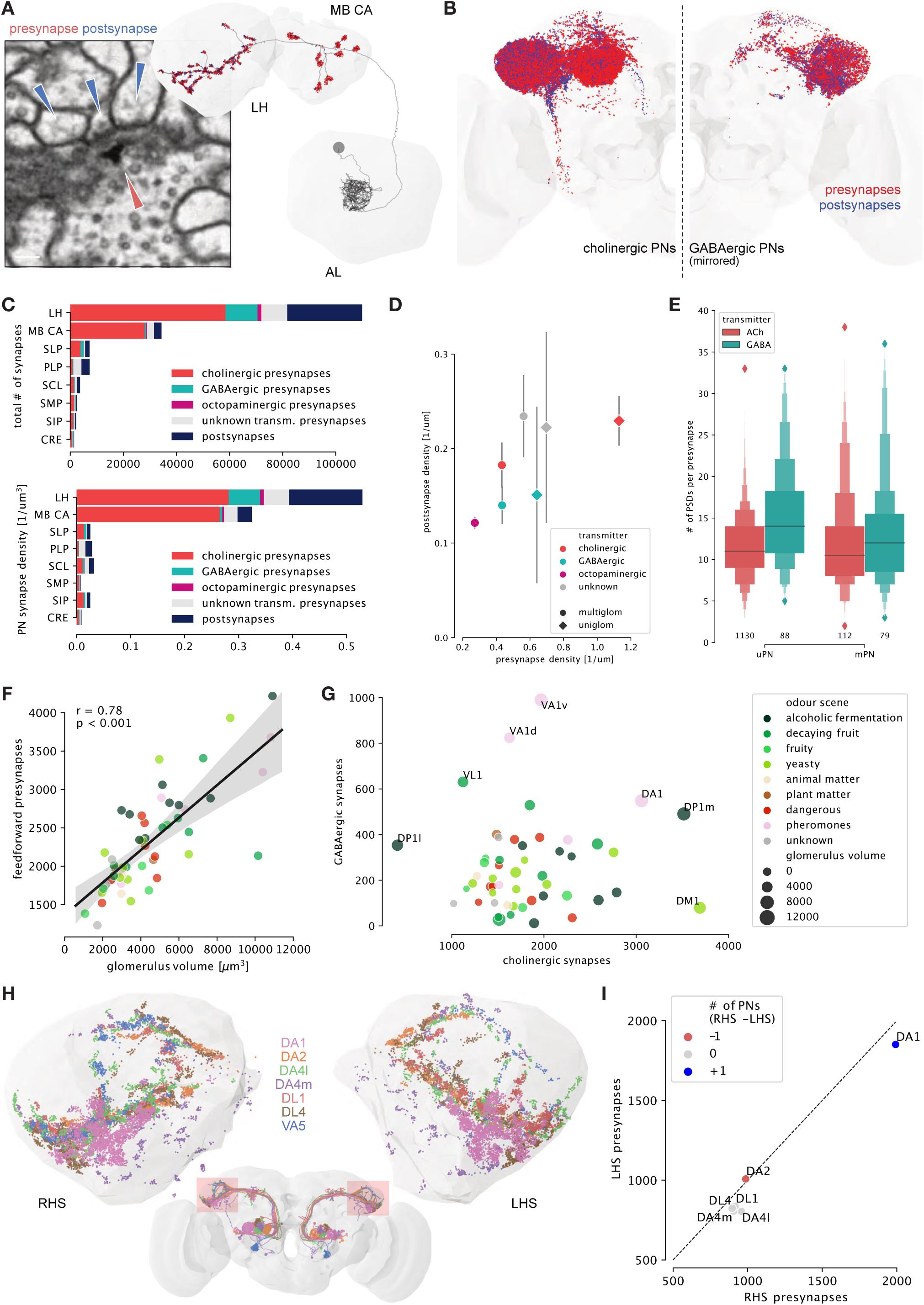
Third-order targets of olfactory feedforward drive. **A** For all olfactory projection neurons (PNs), pre- (red arrow) and post-synapses (blue arrows) were annotated along the axon. **B** Spatial distribution of pre- (red) and postsynapses (blue) by cholinergic and GABAergic PNs (see Figure S2A for Octopamine and unknown transmitter synapses). **C** Total number of synapses (top) and synapse density (bottom) by target neuropil (top 8 shown) and neurotransmitter. LH and CA are the main targets of olfactory feedforward drive (see also Figure S2B). Most modulatory input onto PN axons occurs in the LH. **D** Synapse densities along the axon. All PN types have comparable postsynapse densities. Presynapse densities vary strongly: uniglomerular, cholinergic PNs have more than twice the density of GABAergic PNs. Error bars represent s.e.m. **E** Number of postsynaptic densities (PSDs) for LH presynapses. Bottom numbers represent presynapses randomly sampled to assess synapse sizes. **F** Glomerulus volume strongly correlates with feedforward PN synapses (Pearson correlation, see Methods for details). Colours as in G. **G** Feedforward excitation versus inhibition per glomerulus as function of total number of cholinergic/GABAergic axonic PN presynapses. Glomeruli differ widely in the proportions of excitation to inhibition. Colours correspond to odour scenes. **H** Synapse placement of left/right homologs of 6 uPN types in the LH. **I** Comparison of presynapse numbers between select right- (RHS) and left-hand (LHS) side uPNs summed by glomerulus. Synapse numbers are consistent even in case of mismatches in total number of PNs (see also Figure S2F).

Postsynapse density is fairly constant across PN types (Figure 2D): on average 0.19 ± 0.008 (s.e.m.) per micron of cable. Presynapse density is more variable between PN types: cholinergic uPNs have greater than twice the presynapse density of GABAergic PNs (1.14 ± 0.008 vs 0.47 ± 0.02 per micron of cable). Further, GABAergic presynapses make up only ∼ 15% of olfactory inputs to the LH, which is small compared to the total number of GABAergic PNs targeting the LH.

Presynapses in insects are polyadic [35], i.e. a single presynapse will connect to multiple postsynaptic sites (Figure 2A). For both uPNs versus mPNs and cholinergic versus GABAergic neurons the number of postsynaptic densities per LH presynapse can vary by an order of magnitude but is, on average, very constant across PN types (approximately 12 ± 0.1 s.e.m.) (Figure 2E).

Comparing glomerulus volumes with the number of feed-forward presynapses, we find a strong positive correlation of 0.78 (Pearson R, p <= 0.001) (Figure 2F and S2D, see Methods). Different glomeruli supply different ratios of cholinergic:GABAergic, and uPN:mPN based synapses to higher brain neuropils (Figure 2G and S2E). DM1, for example, spends 86% of its 3898 feed-forward presynapses via cholinergic uPNs, 9% via cholinergic mPNs and only 2% via GABAergic PNs. VL1’s 2404 synapse budget, on the other hand, is spent only to 30% via cholinergic uPNs, 36% via cholinergic mPNs and 26% via GABAergic PNs. This shows that while cholinergic, uniglomerular feedforward excitation is the dominant output of the AL, individual glomeruli can differ.

It is unclear whether the total number of synapses per PN type is similar between brain hemispheres. We chose PN types that consist of a single neuron (DL4, DA4l, DL1, DA4m) and multiple sisters (DA1, DA2), and reconstructed the axons of their left-side homologs to synaptic completion (Figure 2H). The total number of pre- and postsynapses between the left/right homologs are very similar and differ on average only by ∼ 9% (Figure 2I and S2F).

### Axo-axonic connections are widespread between olfactory projection neurons

Reconstructing all right-side PN axons revealed that axoaxonic connections between PN axons are prevalent in the LH and - to a much lesser degree - in the CA, posterior lateral protocerebrum (PLP), superior clamp (SCL) and superior lateral protocerebrum (SLP) (Figure 3A,B). Cholinergic PNs are responsible for most of this PN→PN input (Figure 3D) but receive less PN→PN input themselves than GABAergic PNs (Figure 3C), contrary to prior suggestions [36].

**Figure 3:**
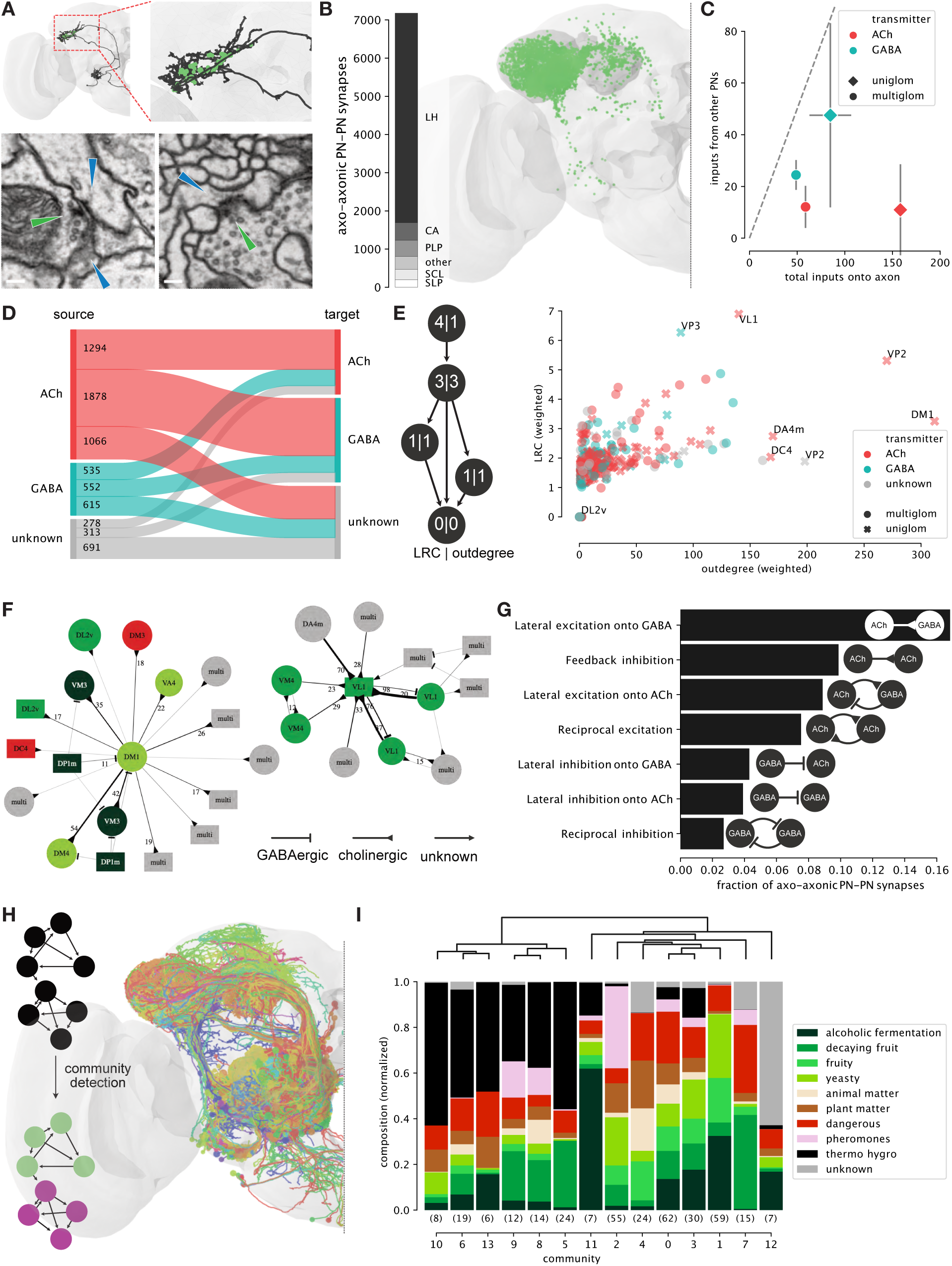
Axo-axonic communities between olfactory projection neurons. **A** Axo-axonic synapses between olfactory projection neurons (PNs) occur onto large backbones (lower left) and small twigs (lower right). Pre- and corresponding postsynapses labeled with green and blue arrows, respectively; scale bar = 100nm. **B** Distribution of axo-axonic PN→PN synapses. The majority of axo-axonic connections between PNs occur in the lateral horn. **C** Fraction of axonic inputs from other PNs. GABAergic PNs receive a higher fraction of inputs from other PNs than cholinergic PNs. Error bars represent s.e.m. **D** Flow chart visualising axo-axonic connections between PNs by neurotransmitter. **E** Hierarchy analysis within the PN→PN network using local reaching centrality (LRC) and outdegree (see Methods for details). A few, mostly cholinergic uniglomerular PNs represent major hubs in the network. **F** Graphs of two exemplary subnetworks surrounding DM1 (left) and GABAergic VL1 (right). Colours as in I; transmitters indicated by arrowheads; numbers indicate synapses per unitary connection. **G** Quantification of motifs in axo-axonic PN-PN network. Arrowheads as in F. **H** Community detection splits PN-PN network into 14 spatially overlapping communities (see Methods). **I** Composition of communities by odour scene shows distinct preferences. See also Figure S3B-C.

In this PN→PN axo-axonic network, 25% of PNs are strongly connected (weighted degree >50) and only 2 PNs (0.6%) are fully disconnected. We find strong connections between sister PNs of the same type (e.g. DA1) but also across PN types, many of which are unidirectional suggesting a hierarchy. We used two metrics to explore hierarchy within the network: the weighted out-degree and local reaching centrality (LRC) [37] (see Methods). Both metrics give similar average results across PN types. Individual PNs appear to be special though (Figure 3E): cholinergic uPNs for DM1, VL1 (both food-related), DA4m, DC4 (both aversive), VP2 (heating), a GABAergic VP3 (cooling) uPN and a few mPNs have above average out-degrees and/or LRCs. DM1, for example, appears to be a hub that connects strongly onto a number of other food-related PNs such as DM4 (Figure 3F). The network exhibits complex connectivity motifs such as feedback inhibition between the excitatory and inhibitory VL1 uPNs (Figure 3F,G). Significantly, the majority of connections are unidirectional - even though in most cases both axons form many postsynapses - violating Peter’s rule and suggesting that some precise developmental tuning mechanism is involved.

The LH can be roughly compartmentalised according to biological valence [5, 38–41]. Seeing that most of the axoaxonic PN→PN synapses occur in the LH, we asked whether we could find a similar organisation within the PN→PN network. We applied a high-fidelity community detection algorithm based on optimising modularity [42] to the PN network. This resulted in 14 large (> 5 PNs) and a few smaller communities (Figure 3H,I; S3C). We then assigned odour scenes (Figure S7) to each community based on the glomeruli innervated by their PNs (Figure 3I) [43, 44]. Community 2, for example, mainly consists of PNs that innervate pheromone-responsive glomeruli DA1, DL3, DC3, VA1v, matching a previously described pheromonal LH compartment (Figure S3C) [38, 39]. Other communities show equally distinct fingerprints suggesting a certain degree of specialisation present in the topology of the network.

### Feedforward olfactory input to neurons of the lateral horn

In order to understand how third-order neurons receive second-order olfactory input, we selected a morphologically diverse set of 66 lateral horn neurons (LHNs) for complete synaptic reconstruction; 40 LH output neurons (LHONs) and 26 local neurons (LHLNs) (see Methods). We identified their developmental lineages of origin [25–28], and their morphological cell type (Supplemental Data, Methods). All examined LHLN cell types are either GABAergic or glutamatergic [15], and therefore likely to inhibit their targets [46, 47]. We contrast these with a set of 15 mushroom body Kenyon cells (KCs) [21].

Most insect neurons comprise a cell body fibre that enters the neuropil and bifurcates, with one branch going on to form its dendrite and the other its axon (Figure S4A). All 66 LHNs, significantly including local neurons, have an identifiable primary branch point and therefore two major arbours (see Methods) (Figure S4A). These can be separated into putative dendrite and axon, based on their density of presynapses (Figure 4B). Some output neurons experience heavy axonic modulation, e.g. 119-228 postsynapses on PD2a1 axons. Others receive very little, e.g. 14-41 on PV5a1 axons. Local neurons also have more presynapses per micron of putative axonic cable than dendritic cable and a slightly greater density of postsynapses on their dendrites. Putative axons are less expansive and thicker than dendrites (Figure S4B). Interestingly, the dendritic compartment is consistently closer to the soma than the axonal compartment (Figure 4B). Together, this suggests that the two compartments are functionally and developmentally distinct in these cells, though it is unclear how they operate electrically.

**Figure 4:**
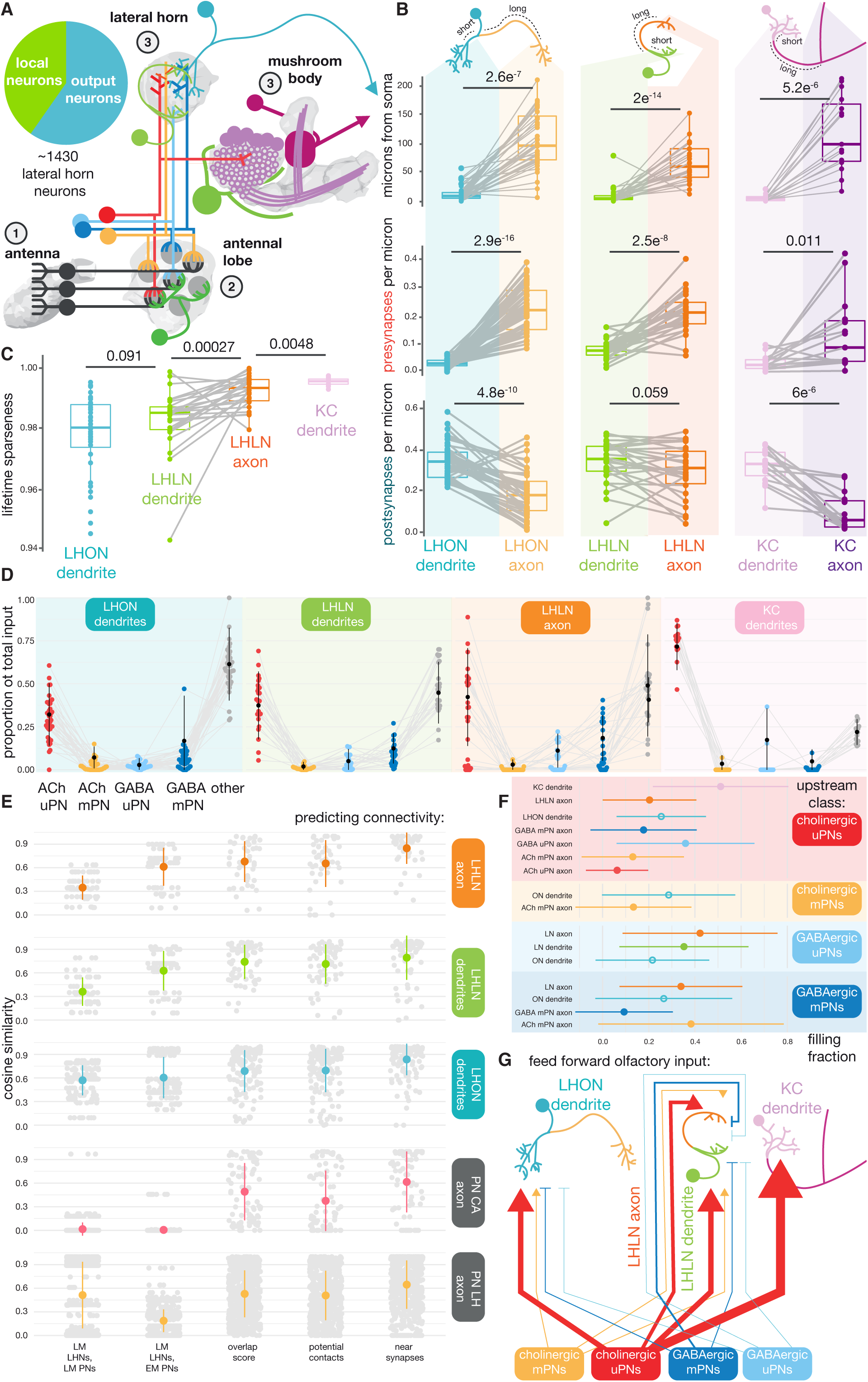
Feedforward olfactory input to innate centre neurons. **A** Neuroanatomical schematic of the first three layers of the insect olfactory system. Pie chart shows what proportion of an estimated 1,400 *D. melanogaster* LHNs per hemisphere, are local or output neurons. **B** Paired boxplots comparing key metrics between for third-order olfactory neurons’ dendrites versus their axons. Paired Student’s T-tests. **C** The lifetime sparseness [45] for third-order olfactory neuron compartments sampling the uPN population (78 cell types). Paired Student’s T-tests. **D** The proportion of postsynapses that are supplied by feedforward olfactory input (colours) from the AL, for all of our reconstructions. Pooled by neuron class and compartment. **E** The cosine similarity between different methods of synaptic prediction and the observed synaptic connectivity from the EM FAFB dataset (see Methods). Only olfactory PN connectivity onto different classes of downstream target (facets) are considered. LM = registered light microscopy data [5, 31]. **F** The mean (± s.d.) filling fraction between two neurons (no. connections predicted by looking for nearby presynapses / observed connections), for olfactory PN connections onto different classes of downstream target. Only results statistically significantly different from the PN→LHON dendrite filling fraction (open circle, blue) are shown. Full results in Figure S4. **G** Schematic of feedforward input from the AL to the LH and CA.

Olfactory PNs account for a mean of 46% ± 14% s.d. of LHNs’ postsynapses, of which 35% is cholinergic (Figure 4D). Each LHN combines excitatory drives from multiple glomeruli largely by combining input from different uPN cell types, which account for 38% ± 15% s.d. of their postsynaptic budget. Of the 78 uPN types, each LHN samples sparsely but not quite as sparsely as KCs. Surprisingly, LHLN axons are sparser than the dendrites of the same cells (Figure 4C). Using a cut-off of 1.5% of total synaptic input, we find that LHONs sample from a mean of 6.9 ± 4.9 s.d. cholinergic uPNs, and LHLNs from 6.0 ± 3.5 s.d. (Figure S4B). This is similar to reports from light microscopy data for both KCs and LHNs [18, 48] (Figure S4B). On average, there is little GABAergic compared with cholinergic or mPN input (Figure 4D), although LHLN axons receive more feedforward inhibition than their dendrites or LHON dendrites.

In order to see whether LHNs selectively wire with certain PNs, we predicted synaptic connectivity between our PNs and 81 third-order neurons through three methods (Figure S4D) (see Methods). Non-synaptic methods tend to over-predict the strength of unitary connections between pairs of neurons (Figure S4B), but even a simple overlap score correlates well with the observed number of synapses, especially in the case of axo-dendritic PN→LHON connections. However, axo-axonic predictions often differed from the observed connectivity (Figure 4E). This is partly because some strong connections are not reciprocated (Figure S5E). For example, DM1, which responds to apple cider vinegar, and DP1m, which responds to the acidic component of vinegar [49], do not axo-axonically connect despite overlapping in space. Instead, they may antagonise one another through PN→LHLN→PN inhibition (Figure S5E). This may allow the fly to better balance the attractive nature of some food sources, against repulsive levels of acidity.

While light-level overlaps can be indicative of some level of connectivity, the innervation densities they suggest across partners may not correlate with our EM data (Figure 4E). This is because the correlation between observed and predicted connectivity for each target neuron varies depending on the neurons’ class and the target compartment. After trying to predict synaptic connectivity based on whether a potential upstream neuron has presynapses near a potential downstream neuron, we find that the proportion to which these unitary different connections are ‘filled’ differs depending on the up- and downstream neuron class and compartment (Figure 4F). For example, cholinergic axo-dendritic uPN→KC synapse creation is far more effective than forming uPN→LHON connections, which in turn are more effective than axo-axonic uPN→LHLN or uPN→cholinergic PN connections. Interestingly, GABAergic PNs are the most effective at forming connections onto LHLNs’ axons.

### Different neuronal classes and compartments have distinct upstream connectomes

Only ∼ 21 % of KC connectivity in the CA comes from neurons other than olfactory PNs. The other elements include non-olfactory projections, a single pan-CA GABAergic interneuron (APL) [51] and other KCs’ dendrites. For LHNs, the unknown slice is larger (Figure 4D): ∼ 58% of LHON dendrites’, ∼ 47% of LHLNs’ postsynapses are supplied by other elements in the LH. ∼ 72% of their upstream connections for PN axons in the LH are not from fellow olfactory PNs. Given that olfactory PNs are the major input to the LH [5], what else could be accounting for all of this synaptic input?

We took all LH PN axons and 7 LHNs (4 LHONs, 3 LHLNs) (Figure S5A), and reconstructed almost all their upstream partners to identification (see Methods). There is a longtailed distribution of synaptic input strengths for the upstream partners of these cells that differs depending on neuron class and compartment (Figure 5A). For example, ∼ 82% of LHLN dendrites and ∼ 86% of LHON dendrites’ upstream partners connect to that dendrite weakly (<1% of the dendrite’s total postsynapses). For axons, this distribution is shifted, with ∼ 15% of LHONs’ and ∼ 33% LHONs’ input synapse budget of incoming connections being ‘weak’, which is more similar to PN axons at ∼ 29%.

**Figure 5:**
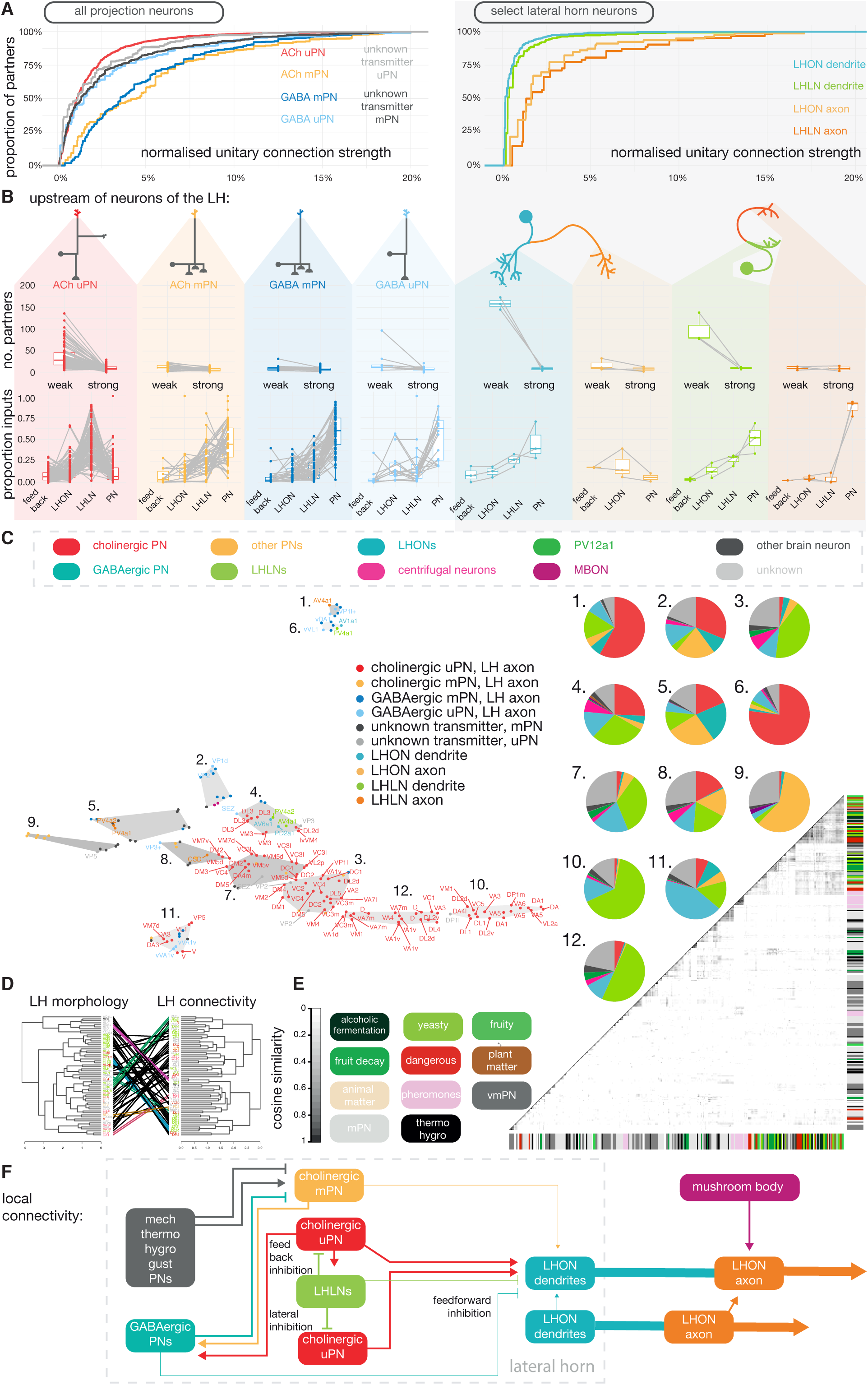
Full upstream connectome for antennal lobe projections in the lateral horn, and for select lateral horn neurons. **A** Left, empirical cumulative density plots for the unitary connection strengths of all upstream partners to downstream targets. Here, the downstream targets are different classes of PN axon in the LH. Right, the classes are different LHN classes and compartments. **B** Upper, the number of synaptic partners that are weak (<1% of target neuron’s synapses accounted for by an upstream partner) and strong (>= 1%). Lower, the proportion of the postsynapse budget for each neuron, that is spent on input from feedback neurons, LHONs, LHLNs and olfactory PNs, where ‘feedback’ refers to input from MBONs and LH centrifugal neurons. Split by class and compartment (coloured facets). **C** A tSNE plot based on synaptic budget vectors for each target neuron, where each value in the vector is the normalised number of synapses supplied by a different upstream neuron class. Grey polygons indicate clusters from a K-means clustering, where k = 12. Pie charts show the how the postsynaptic budget is spent for selected clusters. **D** Tanglegram comparing a clustering based on upstream connectivity similarity (cosine similarity between vectors for synaptic connectivity) and morphological similarity (as assessed by NBLAST [50]). **E** Heatmap showing the cosine similarity scores between the upstream connectivity of individual PNs cell types. Row colours indicate the coarse odour scene in which uPNs are likely to respond [44]. White used for uPNs that respond in multiple odour scenes. **F** Schematic showing the general connectivity motifs this work suggests in the lateral horn.

Local input from LHLNs accounts for about a third of LH neurons’ input (∼ 37% for LHLN dendrites, ∼ 26% for LHON dendrites and ∼ 30% for PN axons) (Figure 5B and S5B). This fraction is only ∼ 4.5% for LHLN axons, suggesting that LHLN axons avoid one another. Both LHON and LHLN input is highly fragmented across many neurons. Only ∼ 10% of LHLN→LHLN unitary connections are strong (>1% of its postsynapse budget), and ∼ 15% of LHLN→LHON unitary connections (a mean of 5 strongly connected LHLNs). However, 84% of LHLN→PN unitary connections are strong (a mean of 7 strongly connected LHLNs).

There is a fair degree of dendro-dendritic input by LHON dendrites, 14% of LHLNs’ and 13% LHONs’ dendritic synaptic budget is spent this way. Even more curiously, LHON dendrites also connect dendro-axonically, accounting for ∼ 5% of LHLN axons’ and 14% of PN axons’ postsynapse budget. GABAergic mPNs supply 18%, a relatively high proportion of LHLN axons’ innervation, given that this figure is <3% for all other categories.

The three LHONs have been implicated in innate attraction [13, 15] and their axons receive a large amount of axoaxonic memory-related ‘feedback’ input (∼ 34%, which is larger than in their dendrites, ∼ 27%). In contrast, AV1a1, a neuron that has been implicated in innate aversive behaviour [17], gets no axo-axonic memory-related feedback, and minimal (0.2%) axo-dendritic. Additionally, ‘attraction’ encoding LHONs receive ∼ 50% of their axonic input from other LHON axons (Figure S5B), while for AV1a1 this is only ∼ 6%.

Distinct fingerprints between neuron classes can be recovered using dimensionality reduction techniques on neurons’ upstream connectivity vectors (Figure 5C). Curiously, some PN axons exhibit more LHN-like fingerprints. Clustering PNs with respect to their upstream partners is comparable to a purely morphological clustering (Figure 5F). Many of these clusters resemble the axo-axonic communities discussed previously (Figure 3I).

We also found that two bilateral, GABAergic PV12a1 [5, 15] interneurons innervate a wide range of PN axons (∼ 5% of all PN axonic input come from just these two cells) and receive input from PN axons (Figure 6D). These neurons may act to normalise activity between the CA and LH, given that each arbourise in both LHs and CAs, and at least in the LH they supply and receive PN connectivity broadly.

**Figure 6:**
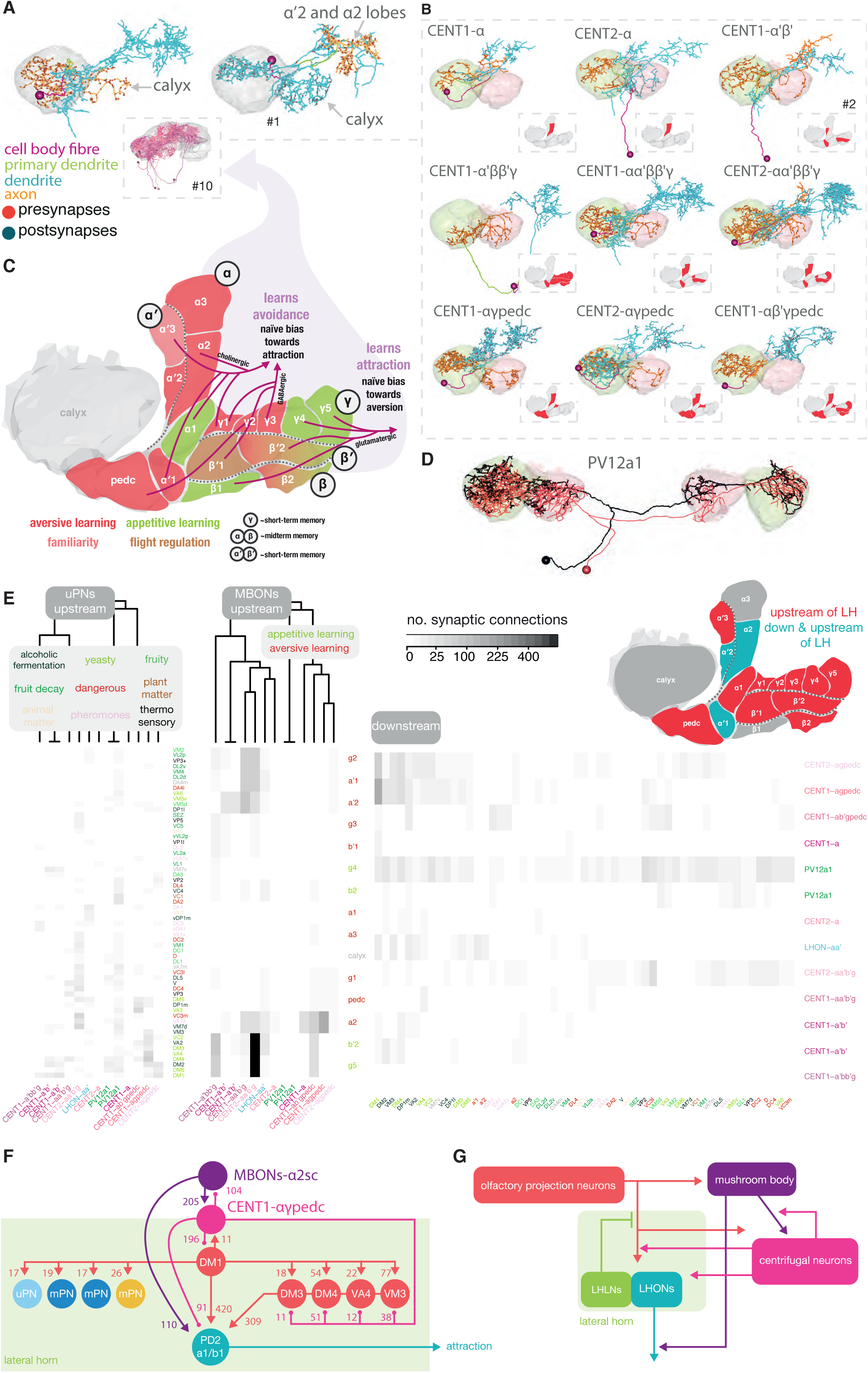
Feedback memory input to the lateral horn. **A** Left, an example of an LH centrifugal neuron. All neurons of the class shown in purple hues, in inset. Right, a single LHON that targets the MB lobes. **B** The 10 LH centrifugal neurons, which form 9 cell types, discovered in this study. All save two also have an axon in the calyx. Insets depict the MB compartments from which MBONs that innervate the LH centrifugal neuron derive. **C** Diagram of the mushroom body lobes, coloured by putative valence [3, 10]. **D** FAFB EM reconstruction of a large bilateral interneuron, PV12a1 [15], that connects both CAs and LHs (black) and its light-level match from a recent split-GAL4 screen (red). **E** Heatmaps showing the connectivity of uPNs onto MB-LH neurons (left), MBONs onto MB-LH neurons by compartment (middle) and MB-LH neurons onto uPN axons and MBON dendrites in MB compartments. PN and MB compartment names coloured by odour scene and valence respectively. **F** Schematic with synaptic weights giving a specific example of how memory can control/modulate control lateral horn neurons known to be important for aversive memory recall [15]. Balls used instead of arrows where the transmitter(s) in use are unknown. **G** Schematic depicting how MBONs interact with LHNs via LH centrifugal neurons.

Curiously, 12% of PN input came from a new class of LH input neuron, putative mechanosensory neurons. Another new class, ‘LH centrifugal neurons’, accounted for 5% (as well as ∼ 11% of LHON and ∼ 2% of LHLN postsynapses) and some of these unitary connections were very high, with the DM1 uPN axon receiving 196 connections from a single LH centrifugal neuron. So what are these neurons?

### Third-order mechanosensory neurons receive innate and learned olfactory information

Previously, we identified four types of mechanosensory GABAergic wedge projection neurons (WED-PNs 1-4) which were predicted to connect to LHONs [15] (Figure S6A,C). Indeed, one LHON, PV5g1#1, receives very little feedforward olfactory input; instead ∼ 10% of its inputs stem from WED-PNs (Figure 4D). We wondered whether WED-PN input may integrate with olfactory information in more prominent ways.

We classified a further four types of WED-PN (see Methods) and fully reconstructed an exemplary neuron for each type (Figure S6C). Unusually, WED-PN 3 and 4 have two dendrites per neuron, one of which is located in the LH (Figure S6A-C). After sampling ∼ 10-70% of the upstream connectivity for exemplary WED-PN 1-4 neurons (see Methods), we found that WED-PNs receive putative feedforward mechanosensory input from the contralateral AMMC and input from local neurons of the wedge. The largest inputs to the LH dendrite of WED-PN 2-4 comes from a class of glutamatergic LHLN, PV4a, and a mushroom body output neuron (MBON) class, MBONs-*α*’3 (Figure S6B). PV4a are classifiers for amine odours [5] that receive uPN input from amine-sensitive and pheromonal glomeruli and whose optogenetic activation induces aversive behaviour [15]. MBONs-*α*’3 are memory-readout neurons involved in olfactory aversive memory recall [10, 52] and the detection of novel odours [53].

We also fully reconstructed MBONs-*α*’3’s cognate dopaminergic neuron (DAN), PPL1 *α*’3 [9]. Interestingly, it receives GABAergic innervation from WED-PNs as well as cholinergic drive from one subtype of these MBONs; MBONs-*α*’3ap but not MBONs-*α*’3m. This motif may be key to implementing their novelty-detection role [53, 54].

### Higher-order brain areas feedback onto the lateral horn

We found that an average of ∼ 2.5% (which can rise as high as ∼ 25%) of inputs to uPN axons is supplied by a hereto undescribed class of neuron. We termed these cells LH ‘centrifugal neurons’ because they feedback to the LH, and often CA, from the LH’s target regions in the superior protocerebrum [5, 15, 18]. We identified and named 10 LH centrifugal neurons as a byproduct of reconstructing PNs’ upstream partners (Figure 6D, Figure S6D). We could not identify a known secondary lineage for most of these neuron. They may be primary cells that persist from the larva. Like MBONs, centrifugal neurons are cytoplasmically large, synapse-dense and therefore metabolically expensive cells. They primarily appear as one neuron per cell type per hemisphere. In the LH and CA, they receive input from olfactory PNs, and also target olfactory PN axons (Figure 6F).

In regions of MBON convergence [9] we observed strong synaptic connectivity from select MBONs (see Methods) onto centrifugal neurons’ dendrites (Figure 6F, Figure S6D); in particular from compartments of the vertical lobe of the MB that play a role in aversive memory and naive attraction (Figure 6E) [53, 55]. In one case where we had a fully reconstructed MBON type, MBONs-*α*2sc, we observed that their axons received LH centrifugal innervation, in this case the CENT1-*αγ*pedc→MBONs-*α*2sc connection was 104 synapses, which is half the count of its reciprocal connection. Centrifugal neurons provide a means by which memory could exercise hierarchical control over ‘innate’ olfactory processing in the LH. However, connections from the LH directly to the site of memory storage, the MB lobes, have not been observed before. However, by following tracts that lead away from the LH we discovered a single example, LHON-*αα*’, suggesting that there is limited LH→MB connectivity but a lot of indirect, and some direct, MB→LH connectivity.

## Discussion

### Numerical stereotypy among olfactory projection neurons

We leveraged a recent synaptic-resolution whole-brain serial section transmission electron micrograph (EM) volume [21] of an adult *Drosophila* brain to take a full inventory of 346 projection neurons (PNs) that relay olfactory information from the AL to higher brain centres (Figure 1C, S1A and Supplemental Data). Because the fly brain is highly morphologically stereotyped, our high-resolution reconstructions should serve as holotypes for identifying these important cells in other datasets [57].

Uniglomerular PNs (uPNs) are morphologically highly stereotyped [38, 58, 59] but genetic driver lines can label different numbers of cells across animals [29]. It is not clear if this represents differences in the underlying cell number or merely variations in expression. A recent study showed a difference in numbers of a single uPN type between left and right hemisphere within an animal [32]. We now tested numerical stereotypy of 58 uPN types across hemispheres and found variability of up to 50% (2 RHS vs 4 LHS VA1d uPNs) in ∼ 17% of cases (Figure 1G).

Interestingly, the brain appears to carefully compensate for mismatches in numbers by adjusting the number of synapses placed by individual PNs. The left/right populations of DA1 and DA2 uPNs each differ by a single PN (7/8 and 6/5, respectively) (Figure 1G). Despite this discrepancy, the total number of axonal pre-/postsynapses is almost identical across hemispheres (Figure 2H,I and S2F). Similar observations have been made for uPN dendrites in one glomerulus [32]. However, PN axons in the LH arbourise in a much more diverse neuropil with a wider range of partners, making it harder to conceive of effective compensation mechanisms. It may be that a PN type’s dendritic postsynapse density and its axonal presynapse density are independently regulated, most likely by their synaptic partners [29, 60]. However, since glomerular size positively correlates with a PN type’s output synapse budget as well as the glomerular OSN count [29], it seems some global mechanism marks out individual PN types to form synapses densely or lightly at both ends.

### Scaling up an olfactory system

The adult olfactory system faces different complex challenges than its larval equivalent. Accordingly, the number of AL glomeruli increases from 21 to 51, the 2.5x increase matching the number of olfactory receptors [33, 61]. The numbers of AL local neurons and uni- and multiglomerular PNs do not scale linearly with the number of glomeruli though but instead increase by factors of 5, 6 and 14, respectively (Figure 1H) [2, 23, 33, 56]. Here, it is striking that mPNs are much more prominent in the fly olfactory system than previously thought. As in the larva, they are downstream of OSNs, other PNs’ dendrites and AL local neurons (Figure S1E). Interestingly, their upstream connectivity is not uniform across glomeruli with some glomeruli providing more feedforward input and other more lateral or feedback connectivity. OSN connectivity is far weaker than in the uPN case [32, 34] and while mPNs are numerous, they possess far fewer output synapses on their axons than uPNs, and so might be less influential as individual feedforward units than uPNs (Figure 2D,E). A non-linear increase in PN diversity may be the product of doubling the number of glomeruli, which exponentially increases the possible combinations that the fly brain could use to decode sensory input.

### Axo-axonic connectivity in the lateral horn

We show that the lateral horn (LH), a region comparable to the mammalian cortical amygdala, is the main target of olfactory projections (Figure 2B-C). Its constituent neurons cover a volume estimated to be 130% of that of the AL [5]; but as with all brain regions, it is limited in space as well as metabolic resources, and keeping the number of neurons and synapses involved to a minimum is critical.

We reveal an extensive hierarchical network of axoaxonic synapses between PNs within the LH (Figure 3). This strongly suggests that some odour channels influence one another, possibly to improve decoding of olfactory information. Implementing axo-axonic connectivity may avoid the need to produce more, metabolically expensive local neurons or additional interneurons with different sampling profiles; enabling economical decoding of evolutionarily-preprogrammed odour valence. For example, the DM1 uniglomerular PN is upstream of multiple, food-odour tuned PNs, such as DM4 (Figure 3F). Since these neurons are cholinergic, it is tempting to imagine that food-related odour channels facilitate one another. However, axoaxonic connections can have non-intuitive effects that depend on the exact timing of action potentials in the connected axons [62, 63].

By connecting to each other in the LH in ways they do neither in the AL nor in the mushroom body (MB) Calyx (CA), PNs enable local interactions that may help the LH categorise odours without affecting the ability of the MB to discriminate odours (Figure 3B). Learning also needs to generalise, but learning outcomes become more general at the level of MB output neurons (MBONs), which can interact directly with ‘innate’ categorisers of the LH (Figure 5B,C) [13, 15].

### Connectivity depends on neuron class and compartment

LHNs have statistically separable dendrites and axons even though all arbours can make and receive chemical synapses (Figure 4B and S4D). Dendrites are consistently located closer to the soma than axons. LHONs have a large number of postsynapses on their dendrites, a mean of 600, and LHLNs have a mean of 545 across both arbours. Upstream of LHNs are hundreds of partners at varying degrees of unitary connection strength; surprisingly, ∼ 80% of LHN postsynapses are spent on weak connections (Figure 5B).

We observe that the filling fraction, the number of predicted synapses that are actually realised, differs between connection types. This hints at class-specific developmental processes that help to create the general motif architecture observed in feedforward connectivity from the AL to the LH (Figure 4F,G). This may happen by using different gains for Peter’s rule [64] depending on the protein expression profile of the neurons involved (Figure 7). For example, individual local neurons can have different input and output relationships with different PN axons, implementing lateral inhibition (Figure S5E,F); similarly PN→PN axo-axonic connectivity is usually not reciprocal. Neuronal activity and/or partner-specific cues may be necessary to specify connectivity after global cues place axons and dendrites in the correct area [65].

**Figure 7:**
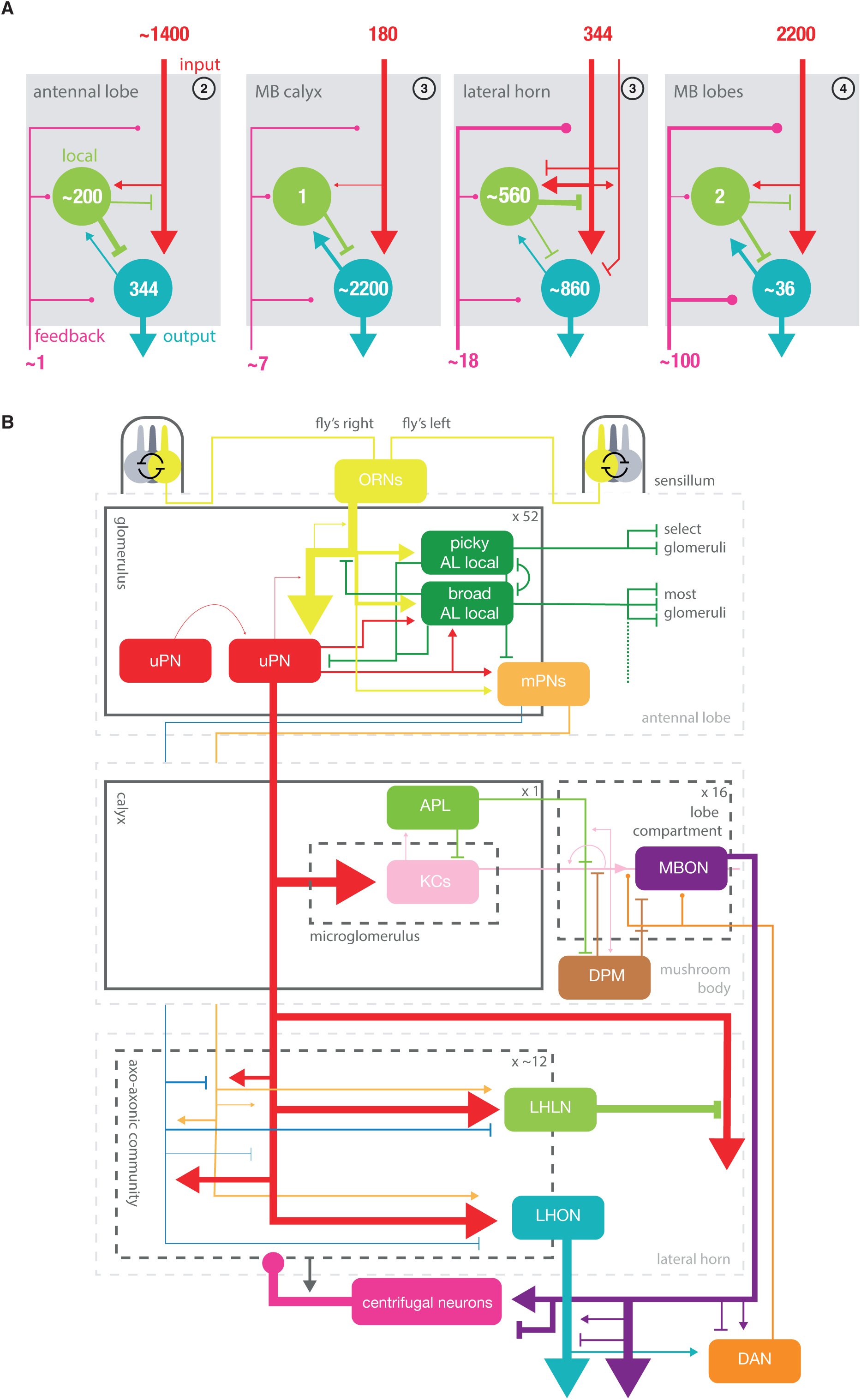
Summary wiring diagrams of the *D. melanogaster* olfactory system. **A** Basic wiring diagrams for four major olfactory neuropils in the fly brain, in terms of input, local, output and feedback neurons. Line widths roughly indicative of the strength of connections, on a neuron:neuron basis. **B** The overall layout of the olfactory system of *D. melanogaster*, including lateral horn wiring uncovered by the present study. 1400 OSNs, 2,200 KCs, 200 AL LNs, 345 PNs, 35 MBONS, 10 MB LH Cent, 1 AL Cent, 100 DANs, 1 APL, 1 DPM, 560 LHLNs, 830 LHONs [5, 9, 29, 48, 51, 56]. Feedback neurons to the AL have not previously been observed (except for the unclearly polarised CSD neurons); we found and partially reconstructed a single, bilateral feedback neuron from the superior protocerebrum to the AL (data not shown).

### Structural differences between olfactory neuropils

Each neuropil contains input, output and local neurons (Figure 7A). The main input neurons to the LH (PNs) are heavily modulated by LH local neurons, making uPN→LHLN→uPN lateral inhibition a prominent feature of the LH (Figure S5C). This is comparable to the situation in the AL and MB lobes where input neurons (OSNs and KCs) likewise receive a significant amount of input from feedback and local neurons [32, 34] -though the numbers of feedback neurons and local neurons are very different (Figure 7A). In contrast PNs also provide axon inputs to the CA but they receive very little modulatory input (Figure 2C). Turning to output neurons, we see that LH output neurons (LHONs) receive a comparatively small fraction (46%) of their inputs directly from olfactory input neurons (PNs). The input profile of LHONs therefore differs markedly from the main output neurons of the AL (i.e. uniglomerular PNs) which receive >80% of their input directly from sensory neurons [32, 34]; they may be more similar to multiglomerular PNs which appear to receive more diverse input (Figure S1E). Conversely, output neurons of the CA (i.e. KCs) resemble uniglomerular PNs in that direct feed-forward excitation accounts for ∼ 80% of their synaptic inputs (Figure 7). Moreover, in the AL, LH and MB lobes the dendrites of output neurons can talk back to input neurons (Figure 5B,C and S1E) while output neurons of the CA (KCs) appear to have few dendritic presynapses (Figure S4D). Among the four neuropils, the LH is unique in that its input neurons (PNs) strongly interact with one another, and that it receives GABAergic as well as cholinergic input (Figure 3). In summary, we see fundamental differences as well as similarities between the first three major olfactory neuropils in the fly brain (Figure 7B).

### Mechanosensory-olfactory integration occurs in the lateral horn

The perception of wind is crucial for animal navigation, particularly for flying animals and for anemotaxis. The LH receives projections from a third-order mechanosensory region of the brain known as the wedge [5, 15, 66–68] (Figure S6A). Activating a line labelling WED-PN1 (R37E08) can generate increased wing-flick motion and differential wing-angling [69] (Figure S6C). We find that WED-PNs receive diverse olfactory input in the LH as well as connections from the the mushroom body from MBON-*α*’3 (Figure S6E). MBON-*α*’3 activation makes flies enter an ‘attentive’ state that suppresses other behaviours [53]. We hypothesise that WED-PNs, as major downstream targets, may mediate this effect. This in turn suggests that WED-PN modulation by olfactory and thermosensory inputs could also result in attention being paid to certain cues, such as polyamines [70], which are of high ethological significance. WED-PNs also target the reinforcing dopaminergic input neuron for the mushroom body *α*’3 compartment. Together, this may represent a circuit (Figure S6E) that directs the fly’s attention in certain contexts or changes its familiarisation-learning rates.

### Memory-related control of ‘innate’ centre neurons

Previously described neurons that directly connect the MB and LH include MB-C1, MBONs-*α*sc, MBONs-*α*’3, PPL1 *α*’3 and PV12a1 [5, 9, 13, 15, 21, 52]. However, we discovered a major new class that we termed ‘LH centrifugal neurons’ (Figure 6B); their dendrites receive input from diverse MBONs (Figure 6E) while their axons targets a range of neurons in the LH, including PN axons (Figure 7). Therefore, multiple MB compartments can influence information processing in the LH. This could be one way in which MBONs directly promote specific behaviours after learning. However the axons of centrifugal neurons in the LH appear to more strongly target PN axons than LHN dendrites; we therefore hypothesise that they may allow the MB to modulate the gain of pathways through the LH rather than directly activating particular behavioural programmes. For example, CENT1-*αγ*pedc is downstream of MBON-*α*2sc and strongly synapses onto the axon of the DM1 uPN (the hub of the food-odour PN→PN community) (Figure S6F). Intriguingly LHON cell types (PD2a1/b1) that receive convergent input from DM1 and MBON-*α*2sc input were recently shown to be required for aversive memory retrieval [13]. The new circuit motif that we describe likely enables the MB to powerfully modify the gain of the attraction-promoting DM1 pathway though the LH. Clearly the MB alpha lobe and LH are strongly interconnected in a way that may give memory hierarchical control over ‘innate’ circuits; this could be used either to suppress those circuits, favouring a learned behavioural response in certain contexts, or to query ‘innate’ circuitry in order to produce desired, learned outcomes. Furthermore, the DM1 PN→PN community could influence the MB alpha lobe via its target, LHON-*αα*’. Possible roles for this connection might be to use food-odours as a training signal or to slow the rate of aversive learning where food is concerned so that the animal cannot easily override its most critical positive instincts. Taken together with other recent work [13–15, 71], we conclude that the MB and LH are much more extensively inter-connected than previously appreciated. Functional studies of this interconnectivity will likely be crucial to a proper understanding of both learned and innate behaviours.

## Supporting information

neuron skeletons

lateral horn neuron cell types

edgelist for lateral horn neuron connections

lateral horn neuron synapses

projection neuron axonic synapses

projection neuron cell types

## Acknowledgements

This work was supported by a Wellcome Trust Collaborative Award (203261/Z/16/Z) to G.S.X.E.J, D.B., G.M.R. and S.W.; an ERC Consolidator grant (649111) and core support from the MRC (MC-U105188491) to G.S.X.E.J; a Boehringer Ingelheim Fonds PhD Fellowship and a Herchel Smith Studentship to A.S.B; a China Scholarship Council award (No. 201908410128) to X.Z. We thank Volker Hartenstein for his help in identifying the lineages that contain lateral horn neurons. Neuronal reconstruction for this project took place in a collaborative CATMAID environment, in which in which 27 labs are participating to build draft connectomes for specific circuits. As a consequence, work done for other published (Dolan *et al.* [13, 15], Zheng *et al.* [21], and Sayin *et al.* [72]) and ongoing projects has also proved useful to us. We thank Fiona Love, Billy Morris, Amelia Edmondson-Stait, Laia Serratosa, Kimberly Meechan, Jawaid Ali, Clement Hallou, Mahmoud Elbahnasawi, Lucia Kmecova, Paavo Huoviala, Zane Mitrevica, Nadiya Sharifi, Istvan Taisz, Irene Varela, Najla Masoodpanah, Melissa Ryan, Peter Gibb, Kaylynn Coates, Shanice Bailey, Quinn Vanderbeck, Jason Polsky, Joseph Hsu, Benjamin Gorko, Jacob Ratliff, Michael Lingelbach, Aidan Smith, Amalia Braun, Adam Heath, Corey Fisher, Kathi Eichler, Ben Koppenhaver, Jeremy Johnson, Przemyslaw Jarzebowski, Remy Tabano, Claire Managan, Johann Schor, Katie Stevens, Adam John, Shada Alghailani, Lindsey Tagg, Philipp Ranft, Bailey Harrison, Steven Calle, Shahrozia Imtiaz, Gabrielle Allred, Markus Pleijzier, Ala Haddad, Emily Tenshaw, Prescott Martin, Austin Warner, Sarah Farris, Georgia Dempsey, Raquel Frances, Dana Galili, Addy Adesina, Levi Helmick, Scott Lauritzen, Farzaan Salman, Andrew Dacks, Kabas Abou Jahjah, Konrad Heinz, Kai Liang and Maria Theiss for together contributing ∼ 30% of the novel neuronal cable used in this study manuscript. We thank Tom Kazimiers for maintaining CATMAID and the CATMAID instances that have served this project. We thank Peter Li for making his automatic segmentation results for the FAFB dataset available to us, and Eric Perlman for making these results available in CATMAID.

## STAR Methods

### 1.1 Neuronal reconstruction

Reconstructions are based on a ssTEM (serial section transmission electron microscope) data set comprising an entire female adult fly brain (FAFB, https://fafb.catmaid.virtualflybrain.org/, http://temca2data.org) (x,y,z resolution 4 nm × 4nm × 40 nm). Generation of this data set was described previously by Zheng *et al.* [21]. Neurons were manually reconstructed using a modified version of CATMAID (http://www.catmaid.org) [73, 74], a Web-based environment for working on large image datasets that has been optimised for tracing and online analysis of neuronal skeletons [74]. Synapses annotated represent fast, chemical synapses matching previously described criteria: thick, dark active zone, presynaptic (T-bars, vesicles) membrane specialisations and a synaptic cleft [75]. We scored each continuous synaptic cleft as a single presynapse regardless of its size or the number of associated T-bars. Adjacent neuronal membranes in contact with the synaptic cleft were defined as being postsynaptic to a given presynaptic site.

PN dendrites were reconstructed to ‘identification’ whereas left-hand-side axons were reconstructed to ‘completion’. A subset (see below) of lateral horn neurons and WED-PNs were reconstructed to completion. MBONs, centrifugal neurons and neurons identified from tracing upstream of completed cells, were reconstructed to identification.

In general, reconstruction to ‘identification’ meant tracing (at least) a neuron’s microtubule-containing backbone in search of major landmarks: in case of PN candidates, for example, we sought to find (a) the soma, (b) axonal projections and (c) dendrites in the antennal lobe in the first pass, such that the neuron’s hemilineage and neuropil-neuropil projections could be discerned.

Reconstruction to ‘completion’ followed the tracing protocol established by Schneider-Mizell *et al.* [74]. In brief, their iterative reconstruction method consists of an initial reconstruction of the entire arbour including annotation of chemical synapses, followed by edits/proofreading by the same or a different tracer. This approach was shown to produce almost no false-positives and to be effective at minimising false-negatives, and has been used by various studies [21, 61, 74].

We modified this protocol for focused proofreading using a recently generated partial automatic segmentation of the data set [76]. In order to help us proofread extant neurons at speed, we semi-automatically looked for missing arbour by using our manual tracings to concatenate disparate, automatically reconstructed neuron meshes, using custom tools we developed (https://github.com/jefferis/fafbseg; https://github.com/flyconnectome/fafbseg-py). Our combined effort to reconstruct the olfactory PNs cost 2180 hours of reconstruction time, 690 hours of arbour review time and created 54 cm of cable, only ∼ 4% of the estimated cable of the LH.

### Choosing neurons to reconstruct

Broadly, there were two approaches when looking for specific individual neurons/cell types: (a) By identifying anatomical loci in the EM that corresponded to anatomical features for our identified lateral horn neurons, for example, their primary neurite tracts. We bridged between loci identified at light-level and the EM in order to build a list of candidate neurons. (b) We used NBLAST to search for our MCFO derived morphologies against extant LH tracing in this dataset. Over the past ∼ 2 years a community of researchers across the world have been reconstructing neurons in FAFB, allowing us to, with their consent, NBLAST against thousands of partial reconstructions to build upon our candidate list for each cell type (see Acknowledgements). Once traced, neurons could be matched to describe light-level data, and so more accurately identified (see below).

We have also attempted to identify the secondary, larval-born lineage of origin for most of our reconstructions, where they were not suspected to be embryo-born primary neurons. There are two naming conventions for a set of about the same ∼ 100 lineage clones, that have originated in the groups of K. Ito and T. Lee, and V. Hartenstein respectively [25–28]. We give both names in our supplemental data, and use the ‘Hartenstein’ names in this text.

#### Antennal lobe projection neurons (AL PNs)

We generated a full catalogue of adult AL PNs by reconstructing all neurons in the three major (mALT, mlALT and lALT) and the heterogeneous transversal (tALT) antennal lobe tracts on the fly’s right brain hemisphere [21–23, 77]. For this, we chose multiple cross sections through the base of each tract and reconstructed all neurons within it to identification. For completeness, we note the serotonergic CSD neuron which we excluded from our list because it does not represent a projection neuron in the classical sense due to its unpolarised nature [78, 79].

#### Lateral horn neurons (LHNs)

The large number of ∼ 1400 lateral horn neurons (LHNs), meant it was not feasible to reconstruct the entire set even to identification. Hence, we chose to reconstruct a sample of 66 to completion. 23 of these have previously been reported [13, 17], and a further 25 correspond to neurons that can be experimentally targeted by specific genetic drivers [15]. 13 were semi-randomly chosen because they exhibited a range of different morphologies. We matched them to 39 defined cell types [5, 57] (Figure S4A) and found four new cell types (supplemental data).

#### LH centrifugal neurons

These neurons were discovered as a by-product of our upstream tracing work (see below). In order to make sure we had as many as possible, we also searched the primary neurite tracts from which these 10 neurons derive, and the heterogeneous tract connecting the LH and MB. We could not identify additional LH centrifugal neurons, though more may exist that do not take these tracts or connect onto PN axons. These neurons were named based on their MBON innervation. If any one MBON innervated an LH centrifugal neuron by at least 4 synapses, the MB lobe in which this MBON has its dendrites was added to the LH centrifugal neuron’s name.

#### Wedge projection neurons (WED-PNs)

First, to find more WED-PNs, we sought to target the tract they take into the LH in FAFB (adjacent to the lALT, ∼ 100 profiles) and the four hemilineages from which they derive (∼ 200 profiles). By tracing these profiles to identification, we identified 23 WED-PNs. All belonged to GABAergic lineages [25]. A further ∼ 10 neurons from the wedge brushed past the LH, including previously studied wind-sensitive WED-PNs [68]. We found that all had an axon in the LH and dendrite in the wedge, with spill-over into other third-order mechanosensory neuropils in the inferior region of the ventrolateral protocerebrum (Figure 6D).

#### Mushroom body output neurons (MBONs)

To identify MBON inputs to LH centrifugal neurons, we transformed segmented neurons from a light-level study into FAFB space [9], and focused on certain MBONs that appeared to overlap with our candidate centrifugal neuron reconstructions (data not shown). After locating the mushroom body compartment and these MBONs’ dendrites, we then targeted their axons for synaptic reconstruction in the vicinity of centrifugal neurons.

#### Upstream tracing from defined neurons

In several cases, we sought to characterise the neurons upstream of cells of interest. In these cases, we reconstructed upstream neurons to identification in order to discover its class [57], where classes included OSNs, uPNs, mPNs, AL local neurons, LHONs, LHLNs, MBONs, KCs, WED-PNs, AMMC local neurons, AMMC projection neurons, PV12a1, LH centrifugal neurons and others. For the reconstruction upstream of 7 fully reconstructed LHNs (Figure S5A), we attempted to reconstruct from all postsynapses, successfully connecting 93% to an upstream neuron that could be classed. To find neurons upstream of mPN dendrites, we sampled fully from one GABAergic mPN dendrite (600 postsynapses) and sampled a random 25% from a larger, cholinergic mPN dendrite (500/2000 postsynapses). We chose random sampling because we were interested in the distribution of synaptic partners across glomeruli. In the case of reconstructing upstream of all olfactory PN arbours in the LH, and WED-PN arbours, we adopted a more efficient sampling strategy to find strongly connected partners, which can be biased by the class identity of that partner [76]. Briefly, this strategy uses a recent partial auto segmentation of the volume to rank potentially connected segments by the numbers of connections they have with the starter neuron, and a human tracer goes through the ranked list. We reconstructed from all such segments predicted to connect by 2 or more synapses with our starter neurons, which focused us away from weak single-synapse unitary connections. Upstream neurons were sometimes assigned a putative modality (e.g. gustatory) if they appear to have dendrites in brain regions known to process particular sensory information (e.g. the subesophageal zone).

### Bridging EM and light-level data

In order to assign glomerular identity to the PNs, we used NBLAST [50] to compare the FAFB EM PNs with segmentations of annotated PNs in the light-level FlyCircuit database ([31, 50]; www.flycircuit.tw). We used a linear, followed by a non-rigid transformation to bring neurons from FAFB into FCWB (fly circuit whole brain) space [50]. Having EM and light-level neurons in the same space, we performed an all-by-all NBLAST and looked for the closest match in the Fly-Circuit database. For the majority of FAFB PNs we were able to find an intuitive match. Identity of PNs types not in the FlyCircuit database (e.g. DP1m) and in cases of ambiguous NBLAST results were manually verified by comparing PN morphology (antennal lobe dendrites, LH arbours and lineage) to published data [57]. We divided the entire EM PN population into uni- and multiglomerular using matches to canonical uPNs as landmarks.

### PN types and putative neurotransmitters

Projection neurons (PN) and lineages thereof have been extensively studied in the past. This allowed us to cross-reference most reconstructed PNs with extant data. In cases of previously unknown PNs, we gave them new type names conforming to the widely used “trivial” names adPN, lPN and vPN (see Figure S1A). In absence of any developmental data, these new trivial names are based entirely on soma position and primary neurite tracts and do not correlate with lineages.

To assign putative neurotransmitters, we assumed that neurons within the same hemi-lineage express the same neurotransmitter(s). This has been shown to be true in the ventral nerve cord [24]. Most extant transmitter data is based on GAL4 driver lines which can have incomplete (i.e. only a subset of a given PN type) or overlapping (i.e. multiple PN types) expression patterns. In addition, individual studies often test only single neurotransmitters and do not show negative staining. We collated this available data and assigned putative neurotransmitters.

In general, neurons contained in the mALT and lALT were shown to be cholinergic based on immuno-histochemical stainings [23]. At the same time, “few if any iACT [mALT] and oACT [lALT] PNs express Gad1” [80]. By itself we did not consider this sufficient evidence for assigning transmitters. Instead we sought additional immuno-histochemical or physiological data:

#### adPN (lineage BAMv3)

PNs in this lineage project through the mALT suggesting they might be cholinergic. In addition, they do not show GABA immuno-reactivity [81]. In electrophysiological experiments neurons in this lineage excite downstream neurons in the lateral horn [18, 20]. We therefore assigned Acetylcholine as putative neurotransmitter.

#### vPN (lineage BAla1)

This lineage contains all PNs of the mlALT and was previously shown to be GABAergic based on immuno-histochemical stainings [82]. Additionally, the mlALT shows no immuno-reactivity for ChAT [22]. Based on this, we assigned GABA as putative neurotransmitter.

#### lPN (hemilineages BAlc ventral + dorsal)

This lineage contains both PNs and AL local neurons. At least some, likely local, neurons are GABAergic [25, 81, 83]. We also know that the PNs in the dorsal lineage hemilineage (contained in GH146-GAL4 expression pattern) are GABA negative [81]. In addition, the dorsal hemilineage contains uPNs that have been shown to excite downstream neurons in the lateral horn [18, 20, 84]. We, therefore, assigned Acetylcholine as putative neurotransmitter to the dorsal hemilineage. Because of the uncertainty with regards to PNs in the ventral hemilinage, we did not assign a neurotransmitter to it.

#### lvPN (lineage BAlp4/ALlv1)

PNs in this lineage project through the mALT and trans-mALTs. It was shown to be neither GABA-, seroton-, dopamin-nor octopaminergic [25]. We therefore assigned Acetylcholine as putative neurotransmitter.

#### VUMa2 PNs

These neurons were shown to be octopaminergic [78]. We note the occurrence of both small clear core and large dense-core vesicles in these neurons, possibly indicative of a second neurotransmitter [85].

#### SEZ PNs: ilPN, ilvPN, ivPN, ivmPN

These PNs originate in the subesophageal zone (SEZ) where they form several distinct clusters. A recent survey of secondary lineages in this region did not identify any that give rise to olfactory PNs [86]. Only a minority of subesophageal lineages divide in the larva to form new adult neurons (28/180), and half have stopped dividing by the late embryonic stage [87]. Therefore, it appears that many small lineages of primary neurons are made in the subesophageal zone, and they likely include some of our PNs, including the VUM cluster. Indeed, similar neurons exist in the larva [33]. We are currently unable to identify exactly which neuroblasts made these PNs. Hence, we assigned new trivial names and included an “i” prefix for “inferior” to indicate their origin in the SEZ.

The biV PNs in the ilPN cluster bilaterally innervate the V glomerulus and were shown to be cholinergic but not GABA-, dopamin- or serotonergic [88]. For other PNs in these clusters we did not find transmitter data.

#### Dorsal PNs: mdPN, dpPN

Although these neurons might well be secondary neurons, we are currently unable to associate them with a lineage. No transmitter was assigned.

### Predicting connectivity

Two methods used only neuron morphology, overlap score and assessment of proximal potential synaptic contact sites, and one method using proximity to presynapses. Connectivity was either predicted using a previously published algorithm for detecting potential synapses [89], or by fitting a linear model to the overlap score versus the numbers of observed connections in EM reconstructions.

In order to quantify the overlap between neuronal skeletons for PNs and LHNs, derived from both light-level and EM data we employed the following ‘overlap score’, as in [5]:

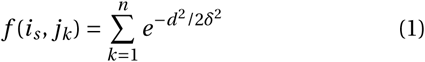

Skeletons were resampled so that we considered ‘points’ in the neuron at 1 *µ*m intervals and an ‘overlap score’ calculated as the sum of *f* (*is*, *jk*) over all points *s* of *i*. Here, *i* is the axonal portion of a neuron, *j* is the dendritic portion of a putative target, *δ* is the distance between two points at which a synapse might occur (e.g. 1 *µ*m), and *d* is the euclidean distance between points s and k. The sum was taken of the scores between each point in i and each point in j.

Overlap scores were calculated between light-level reconstructions from stochastic labelling experiments [15, 31], that have been previously been registered from hundreds of brains to a common template, categorised and identified [5, 50].

To assess ‘proximity’ to presynapses, we took the 90% quantile for distance between connected presynapses and neuron skeleton treenodes (often placed on the neuron’s centreline) (1210 nm). When predicting connections, any synapse from the potential upstream pool (i.e. all olfactory PNs) within this distance to a target neuron, is predicted to connect. 1210 nm was also the value of delta used in the overlap score.

### Compartmentalising neurons

Fully reconstructed neurons were segregated into axon and dendrites using a centrifugal synapse flow centrality algorithm [74], counting polyadic presynapses once. We verified that neurons were suitably polarised by calculating their axon-dendrite segregation index [74] which is a quantification for the degree of segregation of postsynapses and presynapses (0, totally unsegregated, 1, completely polarised). The mean ± s.d. segregation index for PD2a1/b1 neurons was 0.27 ± 0.09 indicating that these neurons are polarised but receive heavy axo-axonic modulation as well as outputting significantly in the lateral horn. MBONs are highly polarised, for example right-side MBONs-*α*2sc had a segregation index of 0.72. Again we counted polyadic presynapses once, rather than using the number of outgoing connections these make, which would have been more expensive in terms of tracing time.

### Morphological Clustering

Morphological analysis using NBLAST [50] was performed on either the dendritic and/or the axonal arbours of neuronal skeletons. Primary neurite tracts and the primary dendrites connecting dendritic and axonal arbours were removed because their fasciculation, especially in the single EM brain space, made NBLAST less sensitive to dendritic and axonal differences. Clustering was performed using functions for hierarchical clustering in base R on euclidean distance matrices of NBLAST scores, employing Ward’s clustering criterion.

### Antennal lobe glomeruli

The antennal lobe (AL) consists of 51 olfactory and 7 non-olfactory glomeruli (see Marin et al., in prep.). Each glomerulus is defined by a specific set of sensory neurons. Because the total number of sensory neurons per AL exceeds 1000, we found it impractical to reconstruct them to define glomerular boundaries [29]. Hence, we opted to use the dendrites of previously described “canonical” uniglomerular PNs (uPN) whenever possible and only reconstruct sensory neurons for glomeruli without any known uPNs. uPN dendrites (and sensory neuron axons) can have extra-glomerular branches which complicates defining cohesive, non-overlapping glomerular compartments. In addition, we found that dendrites of multi-glomerular PNs often appear to be in between glomeruli. We therefore chose a probabilistic approach to calculate an innervation score: we first pruned PNs to their dendrites and generated evenly sampled point clouds. Next, we generated Gaussian kernel density estimates (KDE) for each glomerulus using known uPN dendrites (sensory neurons for glomeruli VP1-5). Using these KDEs, we calculated for each point in a PN’s point cloud the probability (point density function, PDF) for it to be inside of any of the 51+7 glomeruli. Points with a maximum PDF of less than 5e- 15 across all glomeruli were considered to be outside of any glomeruli and hence discarded. For each PN, the final innervation score was calculated by summing up the PDFs per glomeruli and normalised to the sum across all glomeruli. The resulting AL innervation matrix reflects the probability of a given PN’s dendrites being in a given glomerulus.

In order to classify PNs as either uni- or multiglomerular, we set a threshold of an innervation score of >= 0.6 in the top glomerulus.

To calculate the composition of feedforward drive by individual glomeruli (Figures 2F-G, S2D-E, 3I and S3B-C), we assigned each PN’s axonal presynapses proportionally to the glomeruli it innervates by calculating the dot product of the innervation matrix and the number of synapses per PN.

### Network analysis

The weighted out-degree of a node is the sum of the edge weights for edges pointing away from that node.

#### Local reaching centrality (LRC)

The LRC of a node describes the fraction of other nodes that can be reached via outgoing connections [37]. We use the generalisation to directed, weighted graphs that takes the average edge weight (i.e. synapse count) along a path into account:

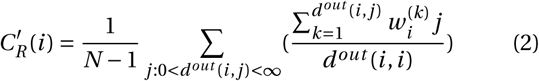

Here *N* denotes the number of nodes in the graph, *d*^*out*^ (*i*, *j*) is the length of the directed path that goes from *i* to *j* via out-going edges and 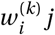 is the weight of the *k*-th edge along this path.

### Tool availability

Morphological and connectivity analyses were performed in R (the natverse, http://natverse.org/) and in Python (pymaid, https://github.com/schlegelp/pymaid) using open source packages maintained by the authors [39]. Figures were generated using Adobe Illustrator, CS6 and Affinity Designer & Publisher. Dataset and source code packages are hosted on GitHub and archived to zenodo.org. All software consists exclusively of open source code released under the GNU Public License and developed at http://github.com.

## Data availability

We provide supplemental data files for the neurons we have focused on in this work. This includes .swc files describing the morphology of each neuron (FAFB14 space) and .csv files with meta data on each neuron (e.g. id number, cell type, lineage), edgelists with the observed number of synapses between neuron compartments and synapse locations in space. Reconstructions will be uploaded to a public CATMAID instance hosted by Virtual Fly Brain (https://fafb.catmaid.virtualflybrain.org/) on publication.

## Supplementary Information

**Figure S1:**
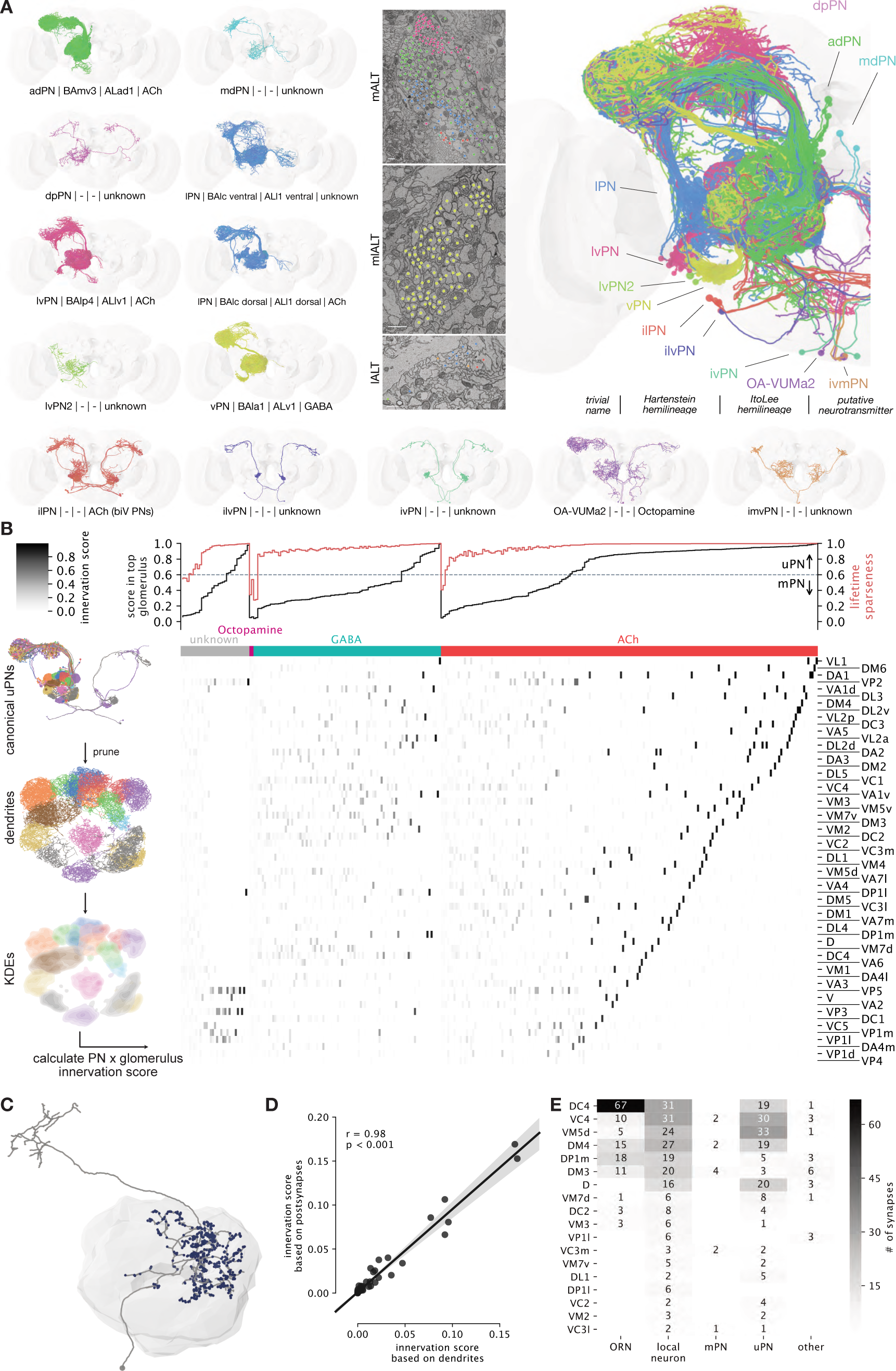
PN lineages, putative transmitters and AL innervation patterns. **A** Projection neurons (PNs) by trivial name; if available, (hemi-) lineage and putative neurotransmitters are given. EM images show cross sections through the three main antennal lobe (AL) tracts (ALTs) with PN profiles highlighted (scale bar 1 micron). Abbreviations: d=dorsal; i=inferior; l=lateral; m=medial; p=posterior; v=ventral. **B** PN by glomerulus innervation matrix based on normalised probability density function (PDF) (see Methods for details). **C** Multiglomerular PN (mPN) which was traced to synapse completion in the AL. Dendritic postsynapses highlighted in blue. **D** Correlation of normalised PDF (i.e. “innervation score”) based on only neural cable and the actual postsynapses of the mPN in C. Strong correlation validates use of neurites to calculate innervation score (Pearson R, p < 0.001). **E** Synaptic inputs to dendrites of mPN in C by glomerulus and neuron type.

**Figure S2:**
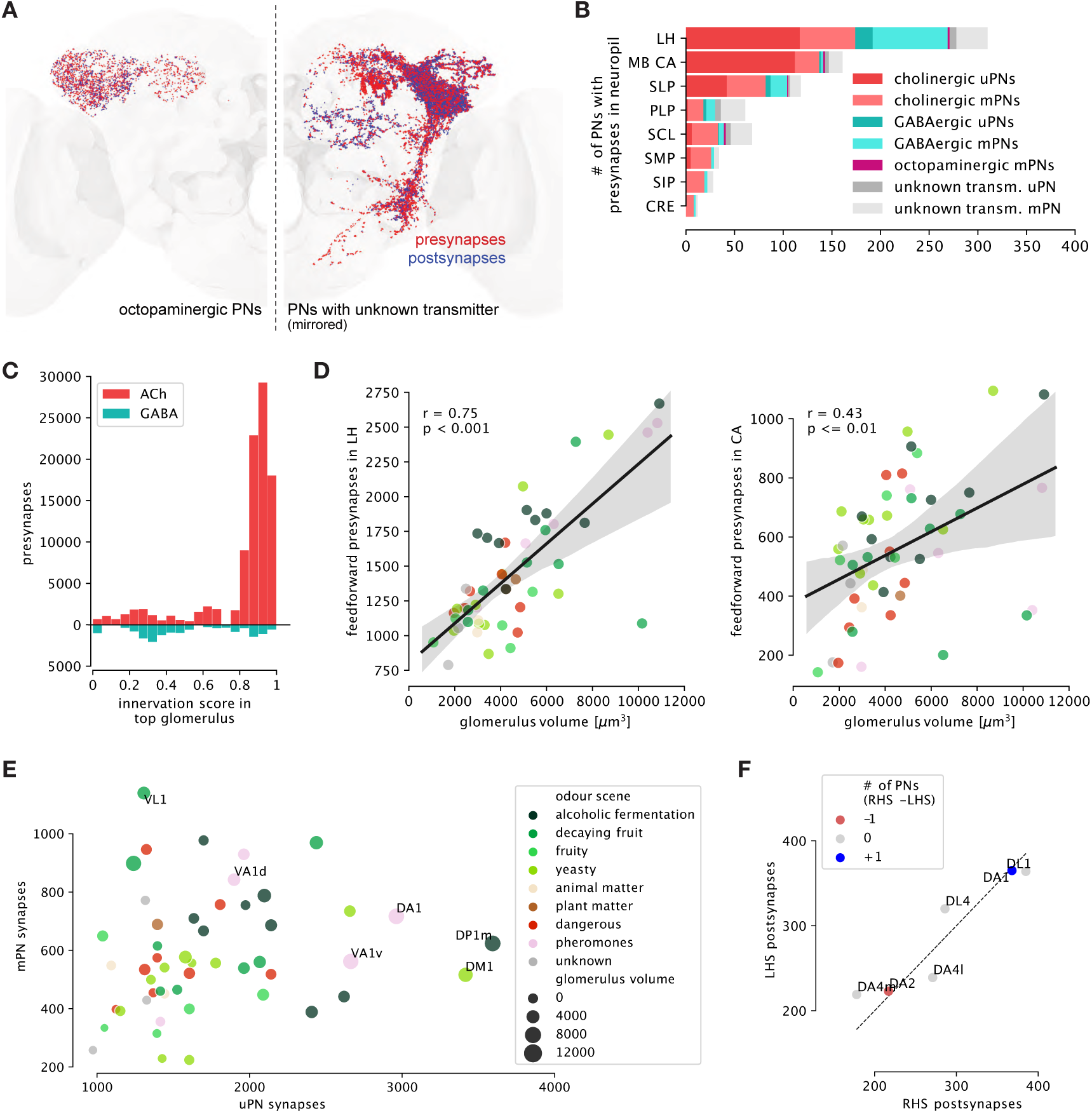
Third-order targets of olfactory feedforward drive - supplemental data. **A** Spatial distribution of pre- (red) and postsynapses (blue) by octopaminergic PNs and PNs with unknown neurotransmitter. **B** Total number of PNs with >10 presynapses in target neuropil (top 8 shown). The LH is the main target of non-cholinergic and multi-glomerular PNs. **C** Cholinergic versus GABAergic feedforward presynapses by sparseness. Sparse PNs make up the majority of cholinergic presynapses while GABAergic presynapses stem from a mix of sparse and broad PNs. **D** Glomerulus volume versus feedforward presynapses in LH (left) and CA (right). Correlation is stronger for the LH than the CA (Pearson correlation, see Methods for details). Colours correspond to odour scenes (Figure S7). **E** Uni- (uPN) versus multiglomerular (mPN) feedforward presynapses vary strongly between glomeruli. Feedforward drive from VL1, for example, comes from equal parts uPNs and mPNs. **F** Postsynapses numbers between left- (LHS) and right-hand-side (RHS) homologs of 6 uPN types.

**Figure S3:**
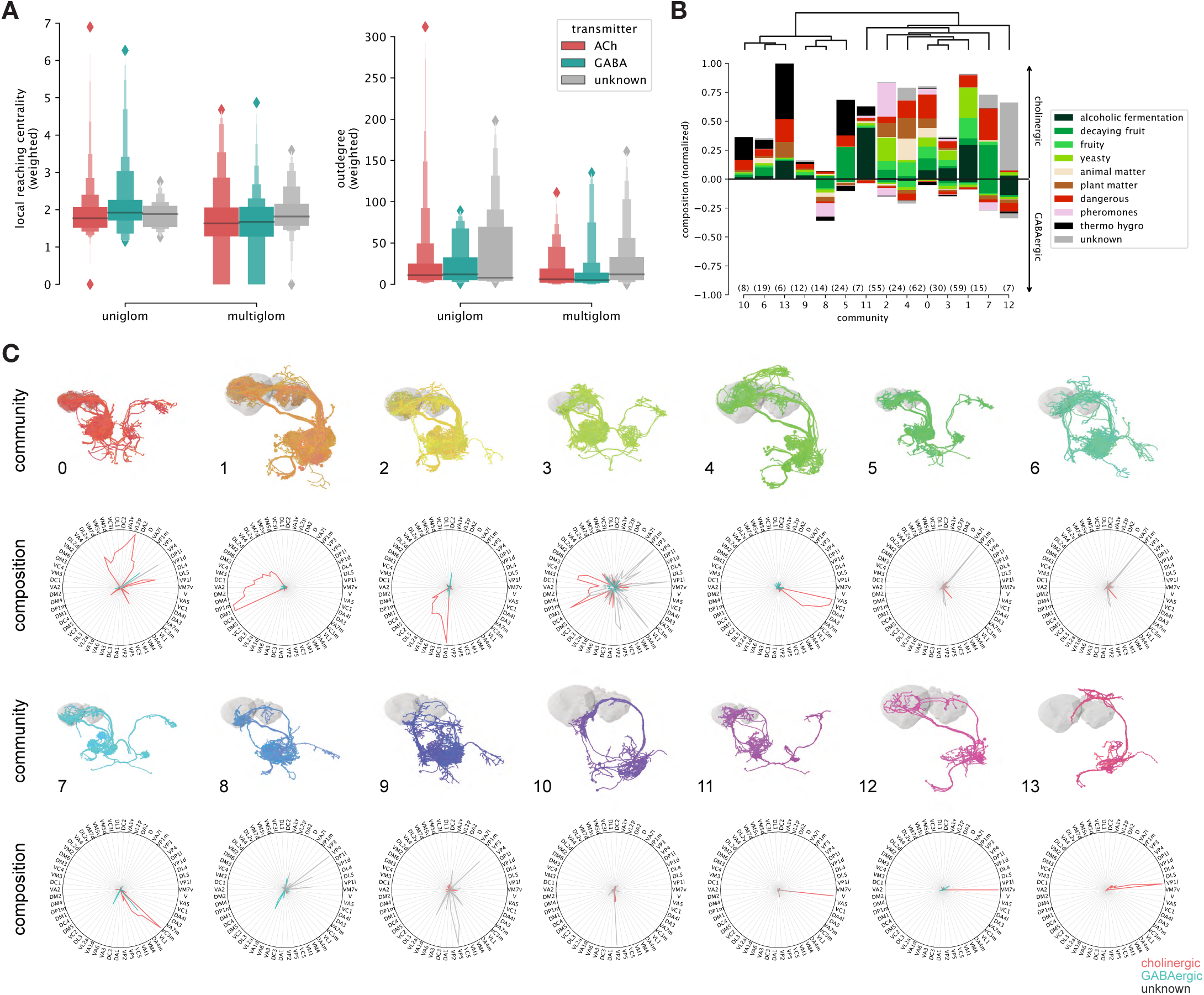
Axo-axonic PN network - supplemental data. **A** Local reaching centrality (LCR, left) and outdegree (right) does not differ between PN types and transmitters. Individual neurons can be strong outliers though (see Figure 3E). **B** Normalised composition of axo-axonic PN-PN communities split by neurotransmitter. The fraction of GABAergic synapses within a community varies between 0% and 35%. **C** Morphology and normalised composition by glomerulus (polar plots) of individual communities.

**Figure S4:**
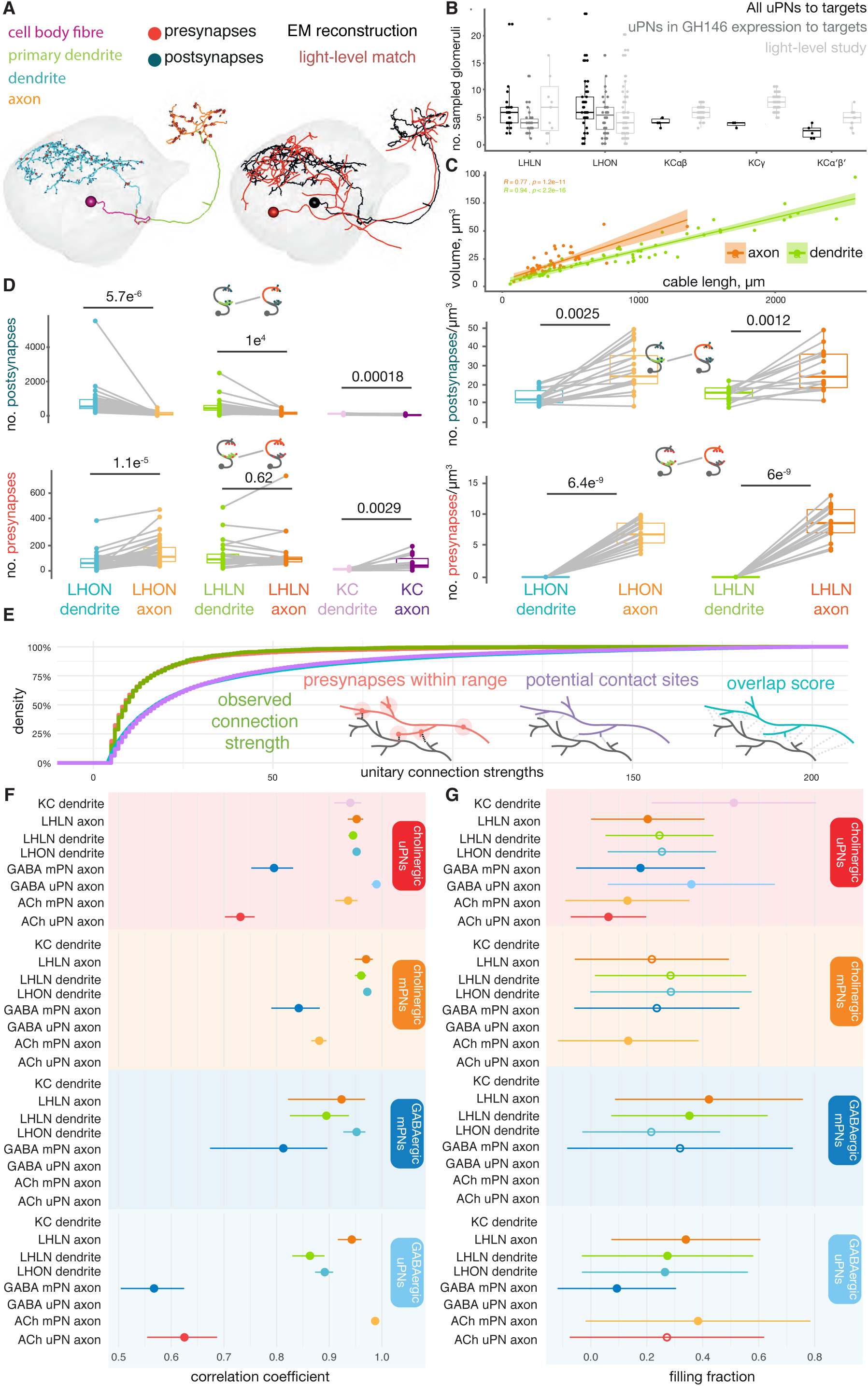
Basic anatomical properties of lateral horn neurons. **A** Left, an example EM complete reconstruction of an LHON, including its synapses and separated neuron compartments [74]. Right, this EM skeleton (black) is matched using NBLAST to a reconstructed skeleton from a library of light-level morphologies (red). This library contains defined cell types [5], and so the EM reconstruction can be identified as AD1a1. **B** Boxplots showing the number of connected glomeruli as assessed from EM reconstructions (black) and light-level studies (light grey). For LHNs, this connectivity comes from a functional assessment of uPN→LHN connections [18], for KCs light microscopy was used [48]. Because Jeanne *et al.* [18] only examined PNs labeled by the driver line GH146, we also restrict our connectivity results to these neurons (dark grey). **C** Correlation between neuronal volume and cable length for dendrites and axons. **D** Paired boxplots comparing key metric between for third-order olfactory neurons’ dendrites versus their axons. Paired Student’s T-tests. **E** An empirical cumulative density plot displaying the unitary (i.e. single neuron to single neuron) predicted and observed connection strengths for reconstructions in our set (66 LHNs). Inset, schematic for three different ways by which synaptic connectivity may be predicted. ‘Overlap score’ and ‘potential contact sites’ do not use any synapse information, ‘presynapses within range’ uses only the position of presynapses. **F** The Pearson correlation coefficients for the relationship between predicted (from the ‘presynapse within synaptic range’ approach) and observed connections between neurons. Closed circles, significant correlations (p > 0.05, Student’s T-test), open circles, non-significant correlations. **G** The filling fraction, the expected proportion of predicted synapses that are realised, is shown. Error bars, standard errors of the mean. Closed circles, significant correlations relative to LHON dendrites (p > 0.05, Student’s T-test), open circles, non-significant correlations. Horizontal, coloured facets indicate the upstream neuron class and compartment, colours indicate downstream neuron class and compartment.

**Figure S5:**
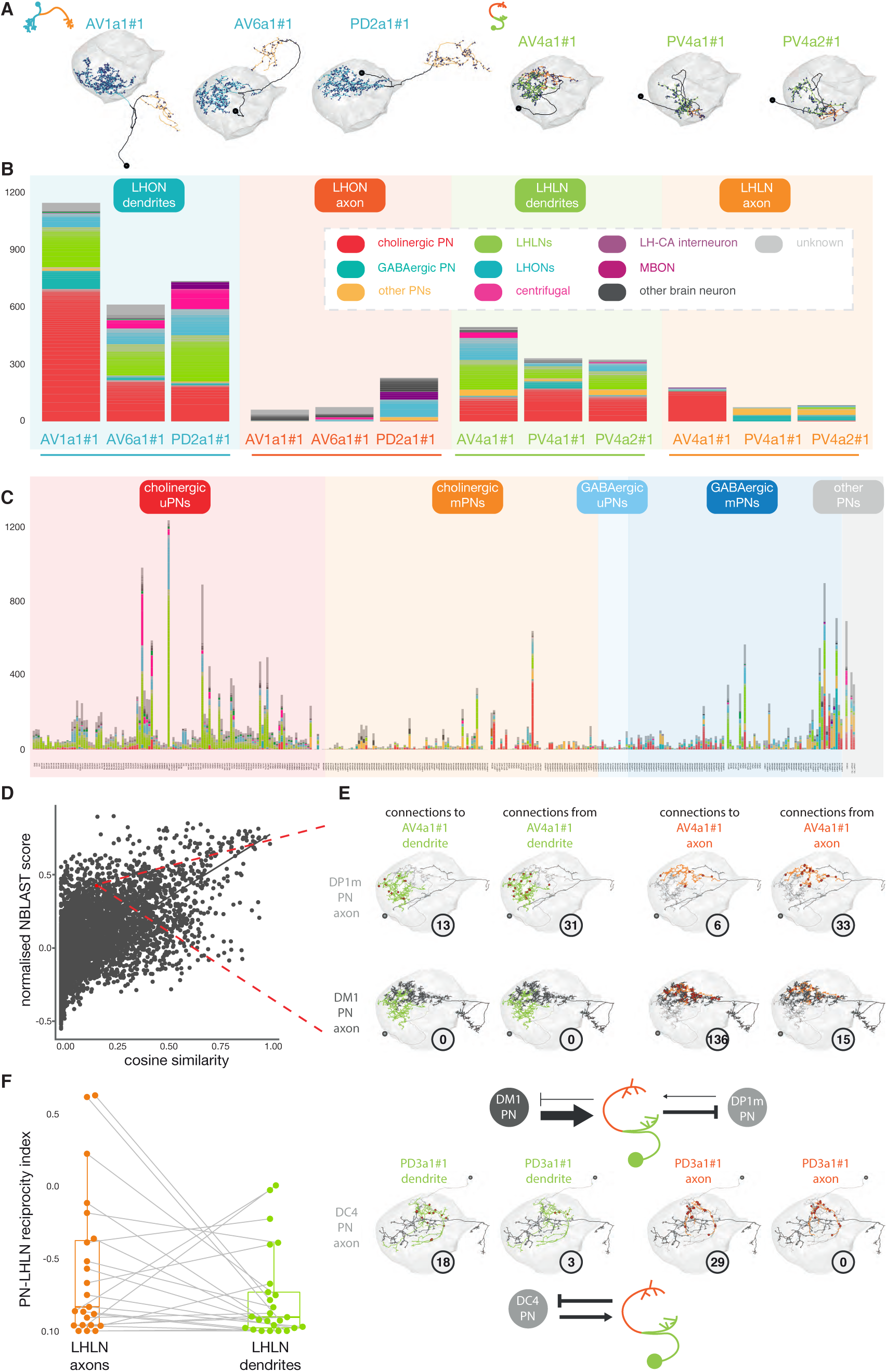
The upstream connectivity of individual neurons in the lateral horn. **A** Left, the three LHONs chosen for full upstream sampling in the LH. Right, chosen LHLNs. **B** Stacked bar plots showing the result of upstream sampling in the LH for these six neurons, faceted by neuron class and compartment. **C** Stacked bar plots showing the result of upstream sampling all PNs in the LH, faceted by neuron class. **D** Scatter plot showing the correlation between PN-PN morphological similarity (NBLAST) and upstream connectivity similarity. **E** Examples of inhibitory connectivity motifs between neurons fully reconstructed in this study, with connection locations shown in red (occasionally obscure one another). Upper, an example of lateral PN-PN inhibition via and LHLN. Lower, an example of feedback inhibition. **F** Paired boxplot comparing the degree to which LHLN axons and dendrites reciprocate connections with PN axons. Plots show an index for connection reciprocity between LHLNs and PNs (LHLN→PN-PN→LHLN connectivity/total).

**Figure S6:**
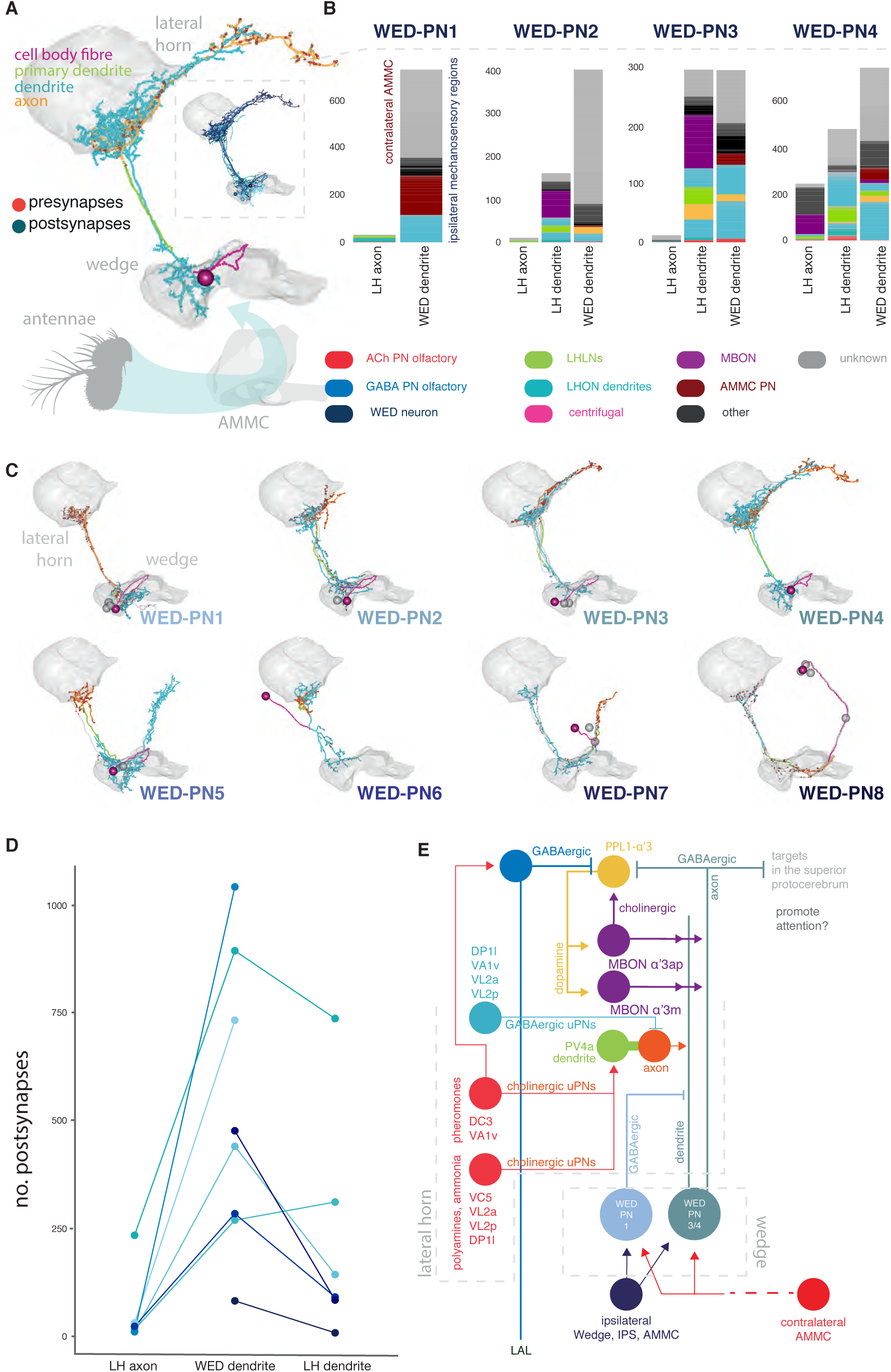
Feedforward wind-sensitive and feedback memory input to the lateral horn. **A** An exemplary synaptic reconstruction of a WED-PN (a WED-PN4) is shown. **B** Stacked bar plots showing the result of upstream tracing from WED-PN compartments, for one exemplary neuron for each of four types. **C** WED-PN reconstructions built from the FAFB dataset, broken down into eight cell types. Exemplary fully reconstructed WED-PN shown in colours. **D** The number of postsynapses different exemplary WED-PNs, broken down by neuropil and neuron compartment. Cell type colours correspond to the colours of labels in C. **E** Putative circuit integrating the results of lifetime learning, via an MBON, with specific, hardwired olfactory and mechanosensory circuitry.

**Figure S7:**
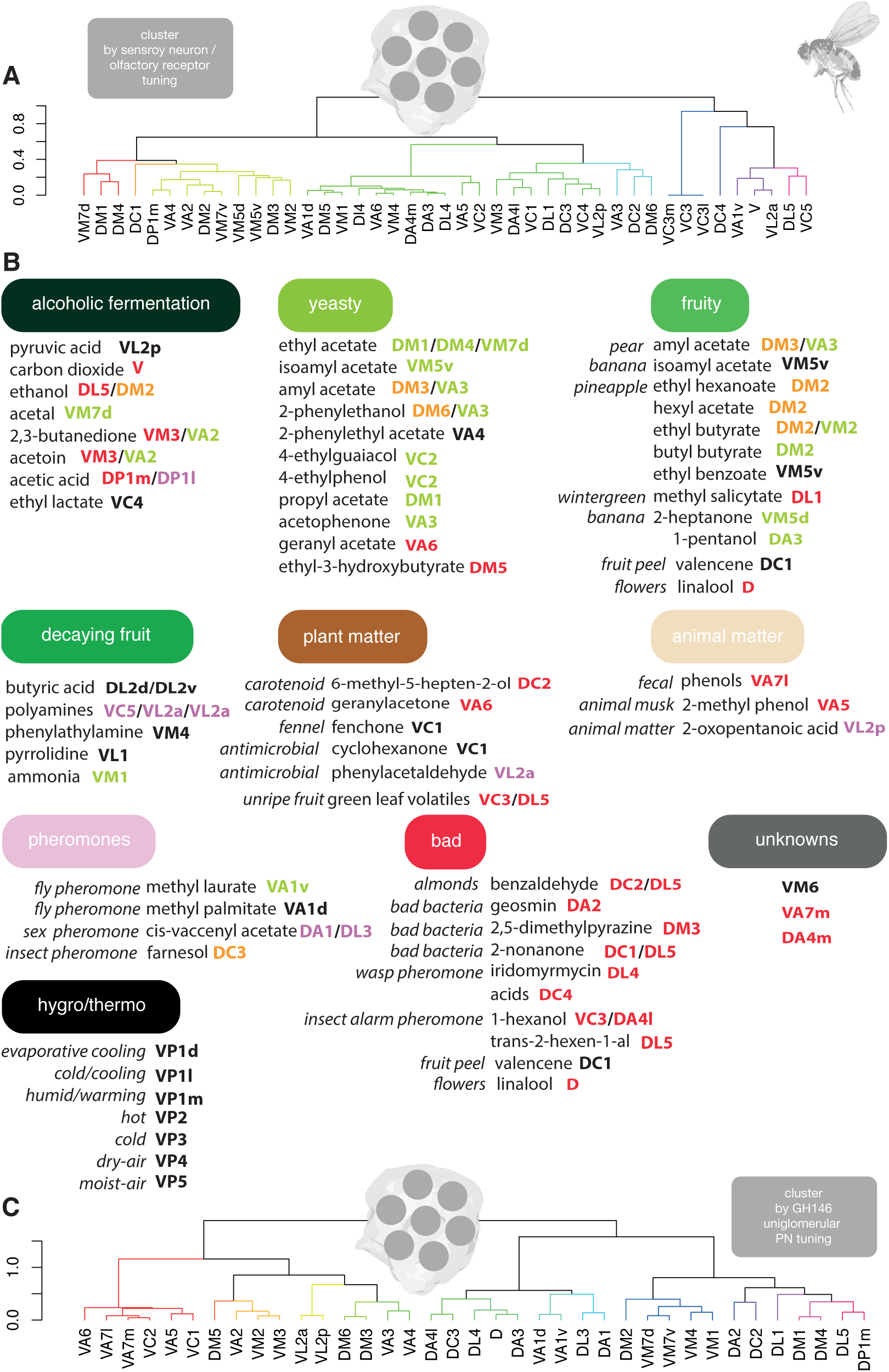
Glomeruli valences and odours scenes. **A** Hierarchical clustering (Ward’s method, arbitrary height cut off for colouring at 0.3) of odour response data from Ca2+ responses in the dendrites of *Drosophila*, uniglomerular GH146 positive PNs [90], after min-max normalisation and calculating their cosine distances. **B** Segregating odour spaces based on a literature review: most prominently using PubChem, the DoOR database, literature summaries from Huoviala *et al.* [17] and Mansourian & Stensmyr [44] as well as other recent work [18, 43, 44, 70, 91, 92]. Example strong ligands for glomeruli are given and in places their occurrence in nature (where this is clear and non-repetitive). Glomeruli are coloured by putative valences (appetitive green, aversive red, black unknown, based on estimates from the literature [43],, purple complex/dimorphic). **C** Hierarchical clustering (Ward’s method, arbitrary height cut off for colouring at 0.3) of manipulated odour response data for ORs and OSNs from the cross-study normalised DoOR 2.0 database. We used the OR to uPN transform described in Olsen & Wilson [93] to estimate uPN responses and then calculated their Gower distances, a method chosen to deal with large amounts of missing data.

